# Organotopic Organization of the Cervical Vagus Nerve

**DOI:** 10.1101/2022.02.24.481810

**Authors:** Nicole Thompson, Enrico Ravagli, Svetlana Mastitskaya, Francesco Iacoviello, Thaleia-Rengina Stathopoulou, Justin Perkins, Paul R. Shearing, Kirill Aristovich, David Holder

## Abstract

Despite detailed characterization of fascicular organization of somatic nerves, the functional anatomy of fascicles evident in human and large mammal cervical vagus nerve is unknown. The vagus nerve is a prime target for intervention in the field of electroceuticals due to its extensive distribution to the heart, larynx, lungs, and abdominal viscera. However, current practice of the approved vagus nerve stimulation (VNS) technique is to stimulate the entire nerve. This produces indiscriminate stimulation of non-targeted effectors and undesired side effects. Selective neuromodulation is now a possibility with a spatially-selective vagal nerve cuff. However, this requires the knowledge of the fascicular organization at the level of cuff placement to inform selectivity of only the desired target organ or function. We imaged function over milliseconds with fast neural electrical impedance tomography and selective stimulation, and found consistent spatially separated regions within the nerve correlating with the three fascicular groups of interest, suggesting organotopy. This was independently verified with structural imaging by tracing anatomical connections from the end organ with microCT and the development of an anatomical map of the vagus nerve. This confirmed organotopic organization. Here we show, for the first time, localized fascicles in the porcine cervical vagus nerve which map to cardiac, pulmonary and recurrent laryngeal function (N=4). These findings pave the way for improved outcomes in VNS as unwanted side effects could be reduced by targeted selective stimulation of identified organ-specific fascicles and the extension of this technique clinically beyond the currently approved disorders to treat heart failure, chronic inflammatory disorders and more.

## 1 Introduction

The functional anatomy of somatic peripheral nerves has been well-studied with serial histological tracing. It has been shown that fascicles observed on a nerve cross-section map reasonably logically to supplied dermatomes and muscle groups. The human vagus nerve is the main peripheral nerve of the autonomic nervous system (ANS) and provides innervation to about eight visceral organs in the thorax and abdomen as well as the larynx. It contains an average of 5 to 8 fascicles but may contain up to 21 fascicles at the cervical level (Verlinden et al., 2016; Hammer et al., 2018a), but, in contrast to the somatic case, their anatomical relation to supplied organs and function is almost entirely unknown (Rea, 2014; Pelot et al., 2020; Ravagli et al., 2020). By homology to the somatic nervous system, it seems reasonable to postulate that fascicles are arranged according to their supply to individual organs and possibly specific functions.

Elucidating the organization of the fascicles in the vagus nerve would be a paradigm shift in the largely unknown functional anatomy of ANS, providing a scientifically advanced understanding of the systems organization of these nerves. This will improve understanding of neurobiological principles and be seminal in assisting studies on neural control (Plachta et al., 2014; Ardell et al., 2015; Bai et al., 2019), neurophysiology, neurological disease and dysfunction (Rajendran et al., 2016; De Ferrari et al., 2017; Asad and Stavrakis, 2019; Drewes, 2021), and ephaptic interactions (Bokil et al., 2001; Capllonch-Juan and Sepulveda, 2020; Sheheitli and Jirsa, 2020). In addition, these findings will be of value in the clinical applications of nerve repair and regeneration (Isabella et al., 2021) and vagus nerve stimulation (VNS) (Rajendran et al., 2019; Thompson et al., 2019; Mastitskaya et al., 2021). Specifically, the latter could be improved with the knowledge of the fascicular anatomy of the vagus nerve by allowing spatial-selectivity thereof and thus avoidance of off-target effects that are frequently experienced such as cough, dyspnea, and bradycardia (Mulders et al., 2015; Fitchett et al., 2021).

Techniques allowing imaging of the anatomy of peripheral nerve *in vivo* include photoacoustic tomography, magnetic resonance imaging (Rangavajla et al., 2014), optical coherence tomography (OCT) (Raphael et al., 2007; Hope et al., 2018; Carolus et al., 2019; Vasudevan et al., 2019) and ultrahigh-frequency and high-resolution ultrasound (Beekman and Visser, 2004; Cartwright et al., 2017; Settell et al., 2021). Unfortunately, none have sufficient tissue contrast, resolution, clarity and penetration depth to trace fascicles confidently along the entire length of the vagus nerve which is >60cm in large animals such as the pig or humans. In addition, these techniques are highly invasive, requiring a large surgical opening to visualize the origin of the fascicles present at the cervical level.

Electrical Impedance Tomography (EIT) is a method that can be used to image and identify organ-specific fascicles within the vagus nerve by correlation of electrical compound action potentials (CAPs) within the nerve with spontaneous rhythmical physiological activity, such as the heartbeat, lung inflation/deflation, or bowel movement (Ravagli et al., 2020). EIT enables the production of images of the internal electrical impedance of a subject using external electrodes. Fast Neural EIT (FN-EIT) enables imaging of neuronal activity within brain or nerve by the detection of small variations in electrical impedance produced by the opening of ion channels during firing and the consequent decrease in membrane resistivity. In peripheral nerves, FN-EIT is performed with a circumferential electrode array set on a nerve cuff, and thus is non-penetrating. It has been demonstrated in rat sciatic nerve with an accuracy of 1 msec and <200μm (Aristovich et al., 2018; Ravagli et al., 2019, 2020, 2021).

Micro-computed tomography (microCT) of peripheral nerve after iodine staining allows *ex vivo* 3D tomographic imaging of the vagus nerve with a spatial resolution of ≈4 μm. This method has been developed and validated for the task of the tracing of fascicles over tens of cm from the innervated organ neural stimulation site to the cervical level in large animals such as the pig or man. This provides independent anatomical validation of any functional connections identified with FN-EIT and selective electrical stimulation (Thompson et al., 2020).

The purpose of this study was to determine the functional anatomy of fascicles in the left cervical vagus of the pig in relation to cardiac, pulmonary, and laryngeal function. The functional anatomy was determined with fast neural EIT as well as trial-and-error selective electrical stimulation using flexible nerve cuffs *in vivo*. These functional measures were validated by *ex vivo* anatomical tracing of fascicles identified in the cervical vagus nerve from their peripheral organ branches, using microCT with iodine staining (Figure 1).

**Figure 1:**
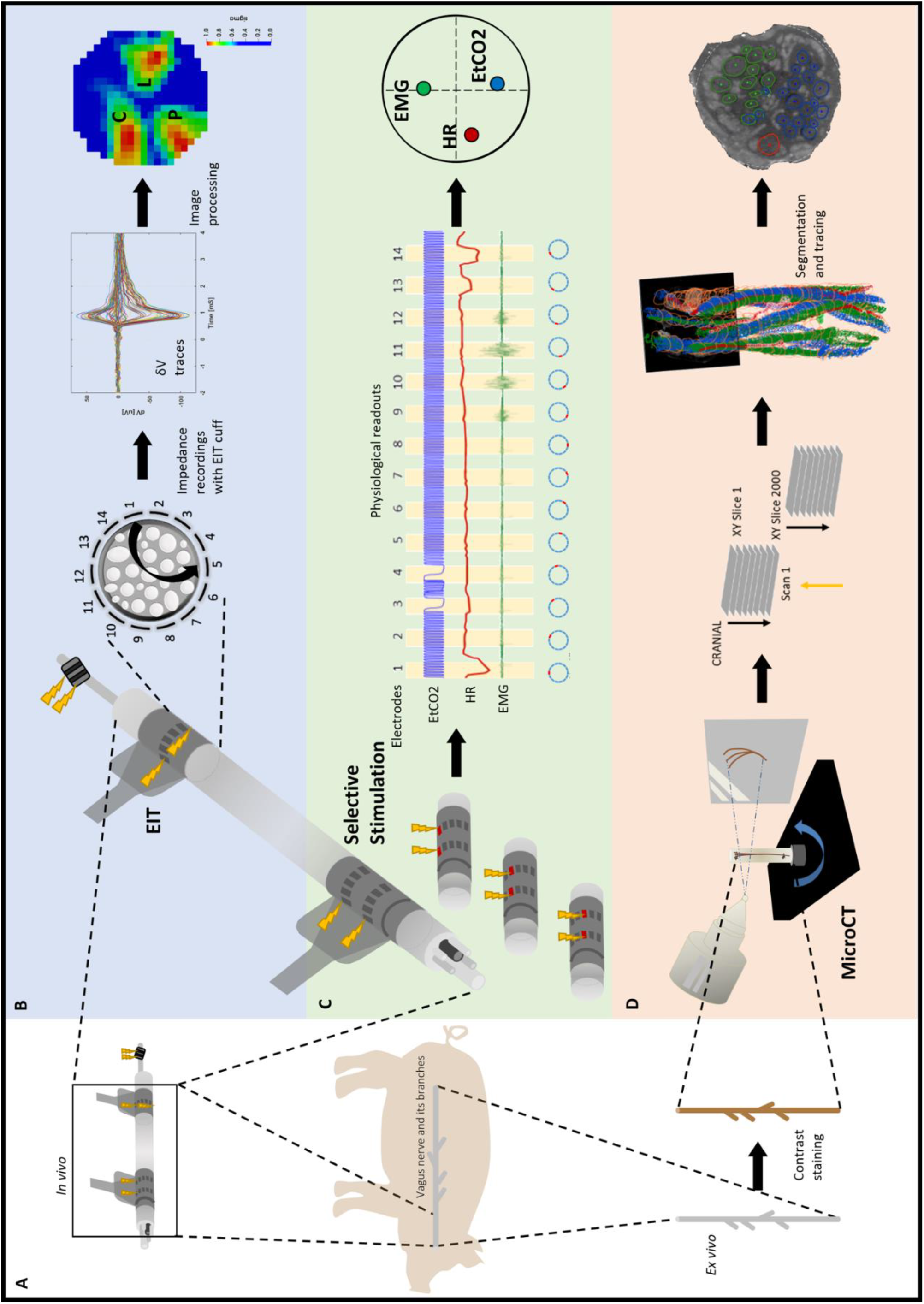
Experimental design for pig cervical vagus nerve imaging with Electrical Impedance Tomography (EIT), selective stimulation (SS), and micro-computed tomography (microCT). (**A**) *in vivo* experiment in pigs with EIT and SS cuffs placed around the left vagus nerve (N=4) followed by dissection of the nerve from cervical level to below the pulmonary branches, including the cardiac and laryngeal branches, for *ex vivo* microCT. (**B**) Identification of the areas responsible for cardiac (C), pulmonary (P) and laryngeal (L) functions in the cervical vagus nerve with EIT. The color scale is arbitrary units (Z score of relative change in the modulus of the impedance). (**C**) Selective stimulation through individual electrode pairs applied sequentially around the circumference of the cuff (pairs 1-14) with resulting physiological changes, such as heart rate (HR, red, cardiac), electromyography (EMG, green, recurrent laryngeal) and end-tidal carbon dioxide (EtCO2, blue, pulmonary), to determine cross-sectional location of the fascicular groups responsible for the respective functions. (**D**) MicroCT scanning of the full dissected vagus nerve followed by segmentation and tracing from the point of organ-specific branching up to the cervical level of cuff placement to identify the location of organ-specific fascicles (cardiac, red; laryngeal, green; and pulmonary, blue).

## 2 Methods

### 2.1 Electrode arrays

Electrode arrays for selective stimulation and EIT imaging were designed to wrap around pig vagus nerves 2.8-3.0 mm in diameter. Electrode arrays were made using the same technology reported in (Chapman et al., 2019) and previously applied to sheep vagus in (Aristovich et al., 2021). Briefly, arrays were made from laser cut stainless steel foil (12.5 μm thick) isolated on both sides with medical grade silicone rubber. There were two designs: 1) EIT arrays comprised one ring of 14 1.50 × 0.35 mm pads and 2 9.00 x 0.47 mm reference ring electrodes placed at extremities of the cuff, shunted to operate imn tripolar mode. Two external reference ring electrodes were present from a template for earlier designs but were unused in this work. 2) Selective stimulation arrays comprised two rings of 14 3.00 × 0.35 mm pads. The design of the electrode arrays used for selective stimulation is also the same as reported and shown in (Chapman et al., 2019) and (Aristovich et al., 2021). Exposed electrode areas were roughened to increase surface area and coated with PEDOT:pTS to reduce contact impedance and noise from the electrode–electrolyte interface (Chapman et al., 2019). This yielded impedances of <1 KΩ and <5°. Cuffs were glued to inner sides of silicone tubing (2.7mm inner, and 4.7mm outer diameter) to maintain tubular shape for wrapping around the nerve.

### 2.2 Acute anaesthetized experiments

The study was performed using Large White domestic female pigs, weighing 60-70 kg. All experimental procedures were ethically reviewed by the UK Home Office and the Animal Welfare and Ethical Review Body (AWERB) and carried out in accordance with Animals (Scientific Procedures) Act 1986. On the day of the experiment, animals were pre-medicated with ketamine (20 mg/Kg) and midazolam (0.5 mg/Kg) administered by intramuscular injection. Fifteen minutes after premedication, a 20 G intravenous catheter was placed in the auricular vein. General anesthesia was induced with propofol (2 mg/kg, i.v.). Animals were intubated with an endotracheal tube, and anesthesia was maintained with sevoflurane vaporized in a 50:50 mixture of oxygen and medical air. A continuous rate infusion of fentanyl (0.2 μg/kg/min) was started after induction and continued during the whole experimental procedure. After induction of general anesthesia, the animal was positioned in dorsal recumbency. Indwelling catheters were percutaneously placed in both the external jugular veins and one in the femoral artery (for blood pressure and blood gas monitoring) using ultrasonographic guidance. The animal was instrumented with ECG leads and a pulse oximeter. A spirometer was connected to the tracheal tube. The animal was mechanically ventilated using pressure control mode for the duration of the surgery and for most of the experiment, except when selective VNS were applied for identification of pulmonary responses. Between these periods, if required, animals were placed onto mechanical ventilation to restore normal levels of CO2 (between 35-45 mmHg). Body temperature was maintained using a hot air warming system if necessary. Ringer lactate fluid therapy at a rate of 5 ml/kg/h was administered intravenously throughout the procedure. Routine anesthesia monitoring included vital parameters such as electrocardiogram and invasive arterial blood pressure, central venous pressure; end-tidal CO2 (EtCO2), end-tidal sevoflurane (EtSev), pulse oximetry and core body temperature (via rectal probe). Some of these parameters (arterial blood pressure, central venous pressure, ECG, EtCO2, EtSev) were also digitally recorded using a 16 channel PowerLab acquisition system (ADInstruments) with LabChart 8 software at 2 kHz sampling frequency. Levels of anesthetic were adjusted accordingly by the anesthetist. In some cases, boluses of propofol or fentanyl were used, if required, and noted on the records. After induction of anesthesia and placement in dorsal recumbency, the ventral neck region was clipped and aseptically prepared using chlorhexidine-based solutions, prior to the placement of sterile drapes, leaving only a small window open for accessing the left cervical vagus nerve. Using aseptic technique, a 20 cm longitudinal skin incision was made using monopolar electrocautery centered immediately to the left of the trachea. The incision was continued through the subcutaneous tissue and the sternohyoideus musculature using a sharp/blunt technique until encountering the carotid sheath and left vagus nerve. A 5-7 cm long segment of the left vagus nerve was circumferentially isolated by blunt dissection to allow placement of SS and EIT electrodes. The electrode cuffs were placed around the nerve by carefully opening the cuffs and sliding the vagus inside it, with the cuff opening facing ventrally (Supplementary Figure 1). The selective stimulation and EIT cuffs were placed at the mid-cervical level, 3 and 5.5 cm from the nodose ganglion, respectively, keeping as consistent placement of the cuffs between animals as possible. To secure the cuff in place around the nerve during the experiment, the sutures incorporated into the design for opening the cuff were tied around the cuff circumference ensuring a tight fit. Electrical ground and earth electrodes were inserted into the surgical field. The impedances of the electrodes were <1 kOhm at 1 kHz. The left recurrent laryngeal nerve was identified within the surgical field, and bipolar stimulating electrode (CorTec GmbH, I.D. 1.2-2 mm) was placed around it. EMG needles were implanted into laryngeal muscle to record laryngeal effects of selective VNS. The surgical field was then rinsed with sterile saline and the skin temporarily closed using towel clamps. After the round of selective VNS for identification of pulmonary fascicles (section 2.4), the animal was put back on mechanical ventilation and anesthesia was switched to α-chloralose (50 mg/kg initial bolus, thereafter 20–35 mg·kg^-1^ ·h^-1^ i.v.). During a stabilization period (30 min) after anesthesia transition, the rounds of selective VNS to identify the recurrent laryngeal fascicles were performed. Selective stimulation for identification of cardiac branches was performed afterwards, and the remaining 2-3 hours of the experiment were dedicated to spontaneous EIT recordings. At the end of the EIT recordings, an overdose of pentobarbital sodium (100 mg/kg i.v.) followed by saturated KCl (1–2 mg/kg i.v.) was used for euthanasia.

### 2.3 Fast neural Electrical Impedance Tomography

Fast Neural EIT imaging to identify laryngeal, cardiac or pulmonary activity was undertaken using a cuff placed 5.5 cm distal to the nodose ganglion at cervical level. The cuff contained 14 electrodes, each 1.50 × 0.35 mm, arranged radially around the long axis of the nerve. EIT images were produced from 196 transfer impedances. Each was obtained by injecting a sinusoidal current with amplitude of 200 μA and frequency of 6 kHz to a pair of electrodes spaced 5 apart (skip-5 configuration) and recording voltages from all 14 electrodes with respect to a circumferential reference ring electrode. The frequency of 6 kHz was identified as optimal for detecting impedance changes in nerves in previous work (Aristovich et al., 2018). The pair of current-injecting electrodes was shifted sequentially to all available 14 radial positions to obtain the 196 traces. Recorded voltage traces were band-pass filtered with a ±1 (larynx) or ±2 kHz (cardiac and pulmonary) bandwidth around the 6 kHz carrier frequency, demodulated with a Hilbert transform and subject to coherent averaging to improve the Signal-to-Noise Ratio (SNR) before being supplied to the algorithm for image reconstruction. Coherent averaging of recorded signals was performed in relation to either triggered activity such as electrical stimulation of the recurrent laryngeal nerve, or rhythmic physiological activity such as the ECG or respiration.

#### 2.3.1 Laryngeal EIT

Laryngeal EIT was performed using a method previously described for the rat sciatic nerve (Aristovich et al., 2018; Ravagli et al., 2019, 2020, 2021). Stimulation was undertaken with a bipolar cuff (CorTec GmbH, Freiburg im Breisgau, Germany) over the recurrent laryngeal branch of the left vagus nerve, approximately 40 cm from the EIT cuff located on the cervical vagus main trunk with biphasic current pulses, 50 μs pulse width and 1.2 mA amplitude at 20 Hz for 60 s for each of the 14 injection pairs (total 14 min). Some δV traces were excluded from reconstruction set due to technical shortcomings, such as poor electrode contact to the nerve or damaged electrode tracks. Traces were excluded if >1-4 μV noise level or 2-6/4-8 μV for mean/max amplitude, which varied slightly for each experiment (Supplemental Table S2).

#### 2.3.2 Spontaneous EIT - Technical

EtCO2, ECG and BP were recorded. Voltage traces were recorded for a duration of 480 s for each of the 14 injection pairs, for a total of 112 min. Demodulated voltage traces (δV) were high-pass filtered with a cut-off frequency of Fc=250 Hz, converted to RMS (root mean square) signals (δV-RMS) with moving average windows of 0.1 s and 2 s for pulmonary and cardiac traces, respectively, and subject to coherent averaging over cardiac- or pulmonary-gated cycles. Coherent averaging is performed in respect to a fixed point of the physiological signal, such as the peak of blood pressure, and allows to improve SNR by averaging the signal over repeated measurements and canceling out the contribution of noise (Aristovich et al., 2018). As for laryngeal recordings, exclusion criteria were applied to identify outlier and artefactual δV-RMS traces and exclude them from the reconstruction set. Exclusion criteria were:

- Trace amplitude larger than 0.2 μV.
- Traces with derivative larger than 50 μV/s.
- Traces deviating more than 3 standard deviations from group average at any given point.
- Traces deviating more than 3 standard deviations from the average Principal Component Analysis (PCA), computed over the group of traces from the same recording and assuming 3 unit vectors.
- Outlier traces characterized a trend over time visibly different from the group ensemble but not falling into the criteria above. Only present in two animals and in negligible amount, 1 out of 144 (0.69%) and 8 out of 144 (5.6%) traces, respectively.

#### 2.3.3 EIT Setup

All evoked and spontaneous EIT recordings were performed using a modified version of the ScouseTom EIT system developed within our group (Avery et al., 2017). Voltage data from the EIT cuff electrodes were sampled in a true parallel configuration at 50 kSamples/s and 24-bit resolution by the EEG amplifier embedded in the EIT system (Actichamp, Brain Products GmbH, Germany), which also contains a 10 kHz hardware filter to prevent aliasing. Modifications to the EIT system performed to improve performance in a surgery room settings included the use of a battery-powered EIT current source (Dowrick et al., 2015) to reduce electrical noise and electrical shielding of the recording system and system-to-cuff cabling.

#### 2.3.4 Image Reconstruction

While spontaneous and evoked EIT recordings underwent different signal processing steps, all post-processed δV traces underwent the same image reconstruction process (Aristovich et al., 2018; Ravagli et al., 2020) (Figure 2). A Jacobian matrix J was obtained from the forward solution (Jehl et al., 2015) to the electrical current distribution problem, computed over a 2.5M-elements tetrahedral model; matrix J was converted into a coarse hexahedral version of voxel size 150 μm and inverted using 0th-order Tikhonov regularization with noise-based voxel correction. For evoked laryngeal EIT, the amplitude of noise-based correction was fixed at 1 μV based on commonly observed noise levels of inter-stimuli signal. For RMS traces resulting from spontaneous pulmonary and cardiac EIT, a fixed noise amplitude level of 10nV was chosen, based on commonly observed levels of δV-RMS signal variability around baseline.

**Figure 2:**
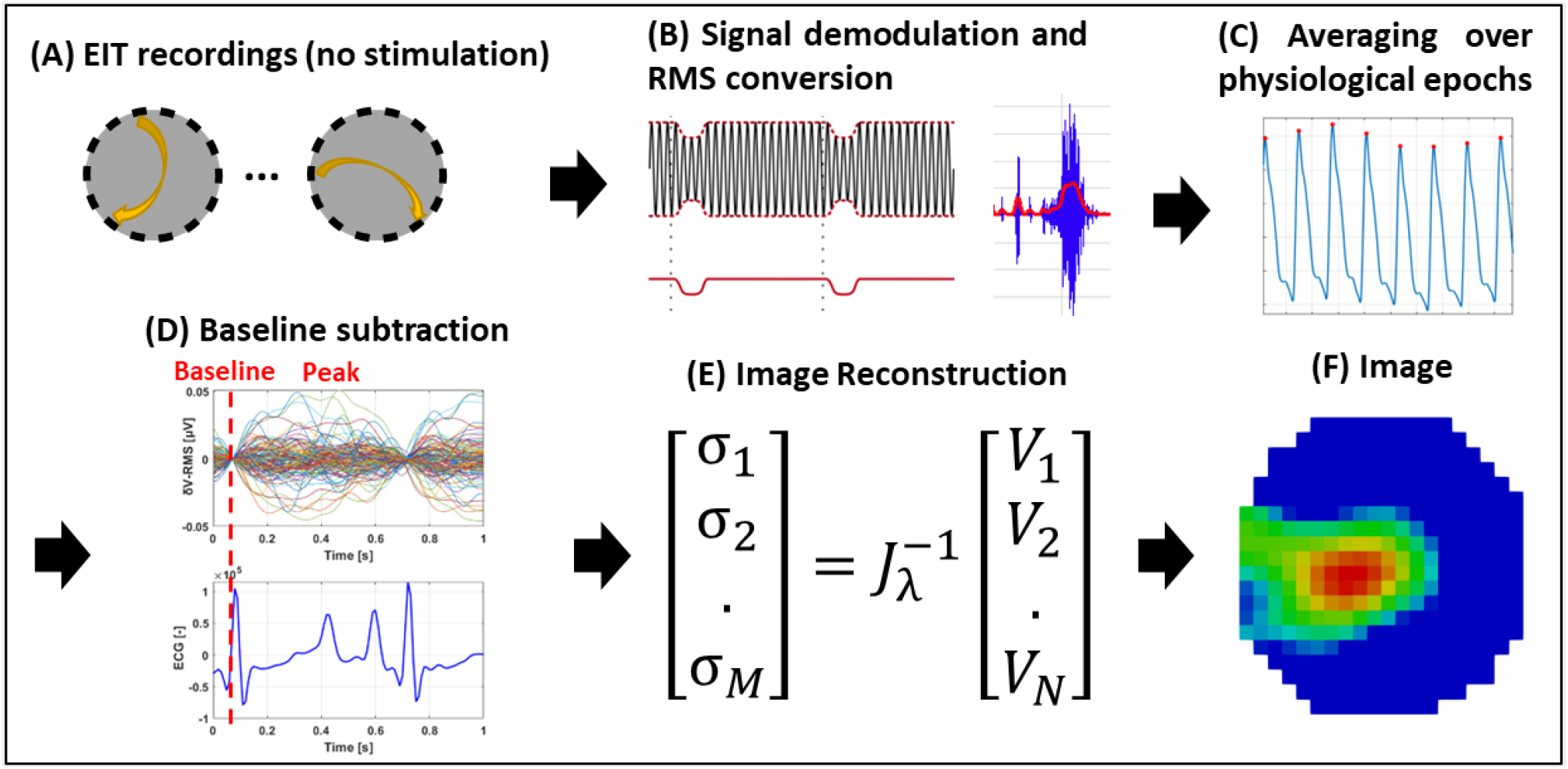
EIT of spontaneous neural activity. **(A)** EIT recordings are performed for multiple injection pairs on the cervical vagus nerve cuff. **(B)** Raw EIT data is demodulated, band-pass filtered and converted to RMS signal (δV-RMS). **(C)** Coherent averaging of RMS-converted EIT data is performed over epochs of periodic physiological signals (e.g., heartbeat or breathing). **(D)** Baseline subtraction is performed over δV-RMS traces at a time point corresponding to low or absent neural traffic. **(E-F)** Tikhonov-based reconstruction is performed to obtain a conductivity map over the cross-section of the nerve. The color scale is arbitrary units (Z score of relative change in the modulus of the impedance).

### 2.4 Selective vagus nerve stimulation

Selective neuromodulation was enabled on a cylindrical cuff with two electrode ring arrays by electrical stimulation through a pair in the same radial position on the two rings and cycling through all 14 available electrode pairs in a consecutive order. The electrodes spaced 4 mm apart around the nerve (Ravagli et al., 2019, 2020; Aristovich et al., 2021). A previous modelling study has demonstrated that this can achieve focused stimulation down to two thirds of the nerve’s radius with an angular spread of 26° (Aristovich et al., 2021). Stimulation was applied in pulse trains at 20 Hz followed by periods of rest of equal duration which were 15 s or 30 s for pulmonary and cardiac or 5 s for laryngeal. For the pulmonary, cardiac, and laryngeal activity, respectively, pulse width (PW) was 50 μs, 1 ms and 50 μs and starting amplitude 400, 1000, and 100 μA. Changes in physiological parameters such as heart rate, breathing frequency, or laryngeal EMG was recorded during stimulation as an index of evoked end organ activity. Pulmonary branch location was assessed by setting ventilation to spontaneous breathing and recording EtCO2 signal as a proxy for breathing rate, while performing selective stimulation on all available electrode pairs. Stimulation of cardiac branch was identified by pair-selective changes in heart rate (HR) measured from systemic blood pressure (BP) or electrocardiogram (ECG). For this purpose, HR drop was computed as the percentage variation between HR at baseline (no stimulation) and during stimulation at each electrode pair. Activation of fibers from the recurrent laryngeal branch was assessed by recording needle-based electromyograms (EMGs) from the larynx during selective stimulation and computing the root-mean-square (RMS) signal (band-pass filtering 5-2000 Hz, RMS window 1 s). Increase of RMS signal from baseline value indicates larynx activation. For each respective organ-specific readout, the stimulation parameters (current amplitude and/or pulse width) were adjusted such that the significant response was elicited on less than 50% of the pairs (less than 7 out of 14). After this, the electrode pair with the maximal physiological response, and all pairs that elicited at least 50% of that response, were selected, and the average center of mass (CoM) angular component for those pairs was computed using their positions around the nerve.

### 2.5 Nerve samples/post-mortem dissection

Following euthanasia, the left vagi of the animals were dissected from the cervical region (with both the stimulating and EIT electrodes attached) down to the branching regions of cardiac, recurrent laryngeal and pulmonary branches, with all the branches left attached to the main trunk of the vagus nerve. All identified branches were dissected and traced back to the originating organ to confirm the suspected, based on gross anatomy knowledge, branch type prior to labeling and cutting the branch (Supplementary Figure 2). At least 1 cm of branch trunk was left protruding from the main vagal trunk from where segmentation of organ-specific fibers would begin. Each sample was approximately 28 cm in length from the upper cervical level (above electrode cuff placement) to beyond the pulmonary branches at the lower thoracic level. Sutures were placed around the vagus nerve prior to any branching region (i.e., the region where a branch leaves the main vagal trunk) as well as to demarcate the positions of the VNS cuffs with the knot in the ventral direction corresponding with the cuff opening. Nerves were then placed in neutral buffered formalin (10%) for fixation.

### 2.6 MicroCT imaging and segmentation

#### 2.6.1 Pre-processing and staining

After fixation, nerve samples (N=4) were measured, sutures of 1 cm length were superglued to the vagal trunk in 4 cm intervals, and nerves cut into 4 cm lengths at the level of suture placement leaving half of the suture on the end of each section as a marker for subsequent co-registration (Figure 3). Two to three sections were placed into a tube of 50 ml Lugol’s solution (total iodine 1%; 0.74% KI, 0.37% I) (Sigma Aldrich L6141) for five days (120 hours) prior to scanning to achieve maximum contrast between fascicles and the rest of the nerve tissue. On the day of the microCT scan, the nerve was removed from the tube and blotted dry on paper towel to remove any excess Lugol’s solution. The nerve sections were placed next to each other onto a piece of cling film (10 cm × 5 cm) (Tesco, United Kingdom) in order from cranial to caudal along the length of the nerve, with cranial ends at the top, and sealed with another piece of cling film to retain moisture during the scan as to avoid shrinkage of the nerve tissue. The sealed nerve samples were rolled around a cylinder of sponge (0.5 cm D × 4.5 cm) and wrapped in another two layers of cling film to form a tightly wound cylinder with a diameter of ~1.5 cm to fit within the field of view at the required resolution. The wrapped cylinder was placed inside a 3D-printed mount filled with sponge around the edges, ensuring a tight fit and the ends sealed with tape (Transpore™, 3M, United Kingdom).

**Figure 3:**
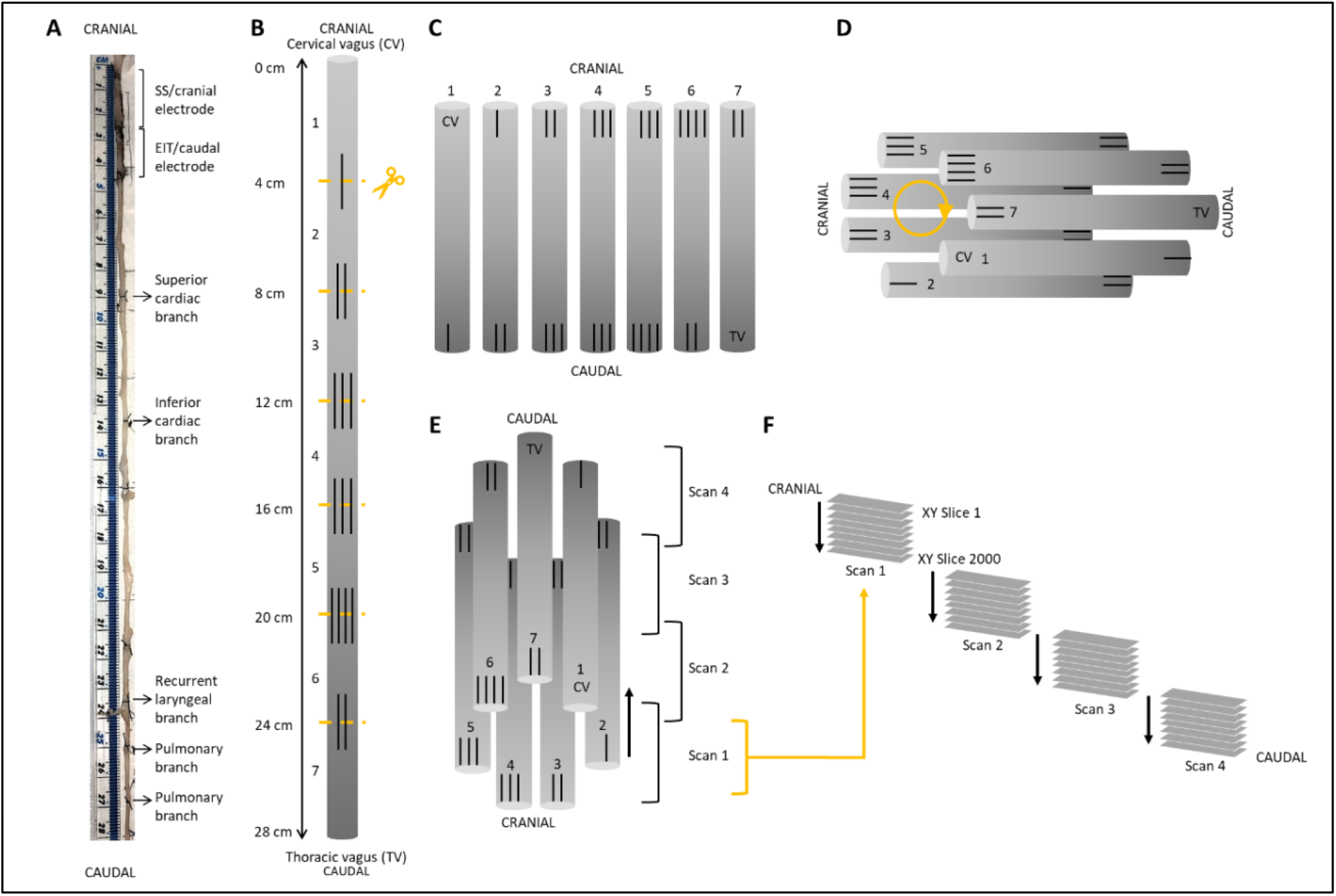
Schematic of nerve preparation. (**A**) Photo of a nerve with sutures tied around the vagal trunk proximal to the branching region and demarcating ends of the two electrode cuffs. Each nerve was approximately 28 cm in length. (**B**) Sutures were superglued into place every 4 cm along the length of the nerve with varying numbers to allow for identification of the sections and alignment during segmentation. Sections were cut at the middle of the markers. (**C**) After contrast staining, nerve sections were ordered from first to last (cervical (CV) to thoracic vagus (TV)), with cranial sides aligned, into cling film and sealed. (**D**) The sealed nerve sections were rolled around a cylindrical sponge in a clockwise order and wrapped in more cling film to maintain its shape. (**E**) The cylinder of nerve samples was placed upside down (cranial side down) into a 3D printed mount and placed in the scanner. (**F**) This setup allowed for the overlapping scans to follow on from another consecutively in the correct order from cranial to caudal along the whole vagus nerve. One nerve is shown here; however, the same preparation was used for all four nerves.

#### 2.6.2 MicroCT scanning and reconstruction

A microCT scanner (Nikon XT H 225, Nikon Metrology, Tring, UK) was homed and then conditioned at 200 kVp for 10 minutes before scanning and the target changed to molybdenum. The scanning parameters were the following: 35 kVp energy, 120 μA current, 7 W power, an exposure of 0.25 fps, optimized projections, and a resolution with isotropic voxel size of 7 μm. Scans were reconstructed in CT Pro 3D (Nikon’s software for reconstructing CT data generated by Nikon Metrology, Tring, UK). Centre of rotation was calculated manually with dual slice selection. Beam hardening correction was performed with a preset of 2 and coefficient of 0.0. The reconstructions were saved as 16-bit volumes and triple TIFF 16-bit image stack files allowing for subsequent image analysis and segmentation in various software.

#### 2.6.3 Image analysis, segmentation and tracing

Reconstructed microCT scan images were analyzed in ImageJ (Schindelin et al., 2012) in the XY plane to view the cross-section of the nerve. The vertical alignment of the nerve was positioned so that the cross-sectional plane was viewed in the XY stack and the longitudinal plane in the XZ and YZ stacks. This allowed for validation of the scanning protocol, direction of the nerve, and visual analysis of the quality of the image and the distinguishability of the soft tissues – specifically the identification of the fascicles known to exist within the nerve. AVI files were created from ImageJ to enable stack slice evaluation, identification of suture positions and branching locations of the vagus nerve, and as a reference during segmentation. Image stacks (XY plane along the Z-axis) were loaded into Neurolucida 360 (Version 2021.1.3, MBF Bioscience LLC, Williston, VT USA) and image histograms adjusted to optimize visualization of the fascicles when required. Fascicles of the three target organs/functions (namely, cardiac, recurrent laryngeal and pulmonary fascicular groups) were segmented from the rest of the nerve using the Contour mode from the Trace tools by forming a closed loop around the boundary of the fascicle of interest. Starting from identification within branches of the vagus nerve, the fascicles were traced through every slice of each scan up the length of the nerve to the cervical region at the level of cuff placement. The suture landmarks placed prior to cutting the nerve into segments were used to match up the neighboring cross sections (Supplementary Figure 3). Contours demarcating the fascicles were placed at regular intervals of 50 to 100 sections and the segmented fascicles labelled accordingly as those coming from an organ-specific branch and thereby containing organ-specific fibers. If fascicles merged with or split into others at a higher frequency, contours were placed at smaller intervals to ensure accurate tracing. While tracing proceeded in the cranial direction, if identified fascicles merged with unlabeled fascicles, the entire new merging fascicle was incorporated and labelled as that fascicular group being traced as it would thereby continue to contain fibers innervating the target organ further up the nerve. If fascicles merged with others already labelled as a fascicular group of interest, it was subsequently labelled as a fascicle containing nerve fibers to both target organs. To continue tracing across cut regions of the nerve, the superglued suture markers and distinct physiological regions or landmarks were used to align the proximal and distal ends of the cut nerves and tracing continued. For visualization of the fully traced nerve, the four overlapping scans were stitched together in Neurolucida 360 by aligning the overlapping regions (approximately 200 slices) of the four sequential scans by removing the artefactual and duplicate sections from the start and end of each scan (100 from beginning and 100 from end), shifting the stack to begin after the preceding scan, and forming one large stack of the segmentation data files. Subsequently, this stack was viewed in the 3D Environment and the contours shelled in 3D.

#### 2.6.4 Histology for validation

Subsequent to scanning, stained nerves were placed back into neutral buffered formalin for a week which allowed for the Lugol’s solution to be soaked out. The formalin was refreshed weekly prior to histology. Nerves were cut into 0.5 cm segments at the level of each cuff placement. The segment was embedded in paraffin, sectioned at 4 μm, stained with Hematoxylin and Eosin (H&E, a routine stain used to demonstrate the general morphology of tissue) (Sheehan and Hrapchak, 1987), and imaged with light microscopy. Identification of histopathological features was performed, and the images of the H&E sections were then compared to the corresponding slice in the microCT scan of the same nerve for comparison and validation. The presence and number of fascicles visualized in the golden standard of histology was compared to those identified during segmentation of microCT scans to confirm that what was segmented was correct.

#### 2.6.5 Nerve analysis

In order to calculate the difference in organization and positioning at the cervical level, the following was determined. The order and location of the three fascicular groups of interest within the nerve were analyzed for each nerve and compared between samples; the cuff opening was used as a reference point, being located ventrally. The number of fascicles present in each of the vagus nerves at the level of both cranial and caudal cuff placements was counted for the vagus as a whole, and within each of the fascicular groups namely cardiac, recurrent laryngeal and pulmonary, as well as for the number of ‘overlapping’ fascicles between groups. The diameter and area of the whole vagus nerve was calculated from the histology images using ImageJ Analyze-Measure function after both the scale was set and the borders of the nerve were segmented. The area of each of the fascicular group regions identified, the overlapping regions, and the total area containing fascicles was calculated as a percentage of the area of the whole nerve from the microCT images. The distance between the cervical level (cuff placement) to the level of branching was calculated for each branch of the three groups/functions. Full morphometric analyses of the nerves was performed at the level of the two cuff placements and of each branch (see Supplementary Files). All counts and measurements were compared between nerves.

### 2.7 Image co-registration, quantitative metrics and statistical analysis

SNR was computed for EIT recordings as the ratio between average δV (for laryngeal branch EIT) or δV-RMS (for pulmonary/cardiac branches) signal at peak variation and baseline/pre-stimulus average noise. The neuromodulation effect for each branch was computed as variation from baseline value evaluated on the most effective electrode pair defined as having the greatest change in a physiological readout correlating with the organ-specific function. Those electrode pair(s) that induced more than 50% of the maximal physiological response during selective stimulation were chosen to compute the angular component of the CoM for this technique; radial component was chosen as two thirds of nerve radius. The angular distance of pulmonary and cardiac activation peaks from laryngeal were computed for EIT, SS and MicroCT. Discrimination power was assessed by T-test for each technique. Spatial precision of selective stimulation and localization power of EIT imaging were assessed by comparing CoM locations from the two techniques with CoMs identified from microCT imaging at each cuff. For EIT images, coordinates of peak spontaneous or evoked neural traffic were identified by computing the CoM of the highest-intensity 16 voxels (4×4 area) over the cross-section of the nerve at the time of signal peak. For selective stimulation, focal points of activation for each branch were identified by computing the average polar coordinates of all electrode pairs inducing clearly observable significant neuromodulation. Selective stimulation currently offers only angular steering capabilities and no depth resolution, so radial coordinate was fixed at 2/3^rd^ of the radius, corresponding to 1mm from the nerve center, out of a 1.5mm radius. This was based on the previous sVNS study indicating that sVNS activates fibers in the outer part of the nerve (Aristovich et al., 2021). The angular coordinate was computed as the average of angular coordinates of the electrodes inducing neuromodulation effects during selective stimulation, such as electrodes 8-9-10 in Figure 4.

**Figure 4:**
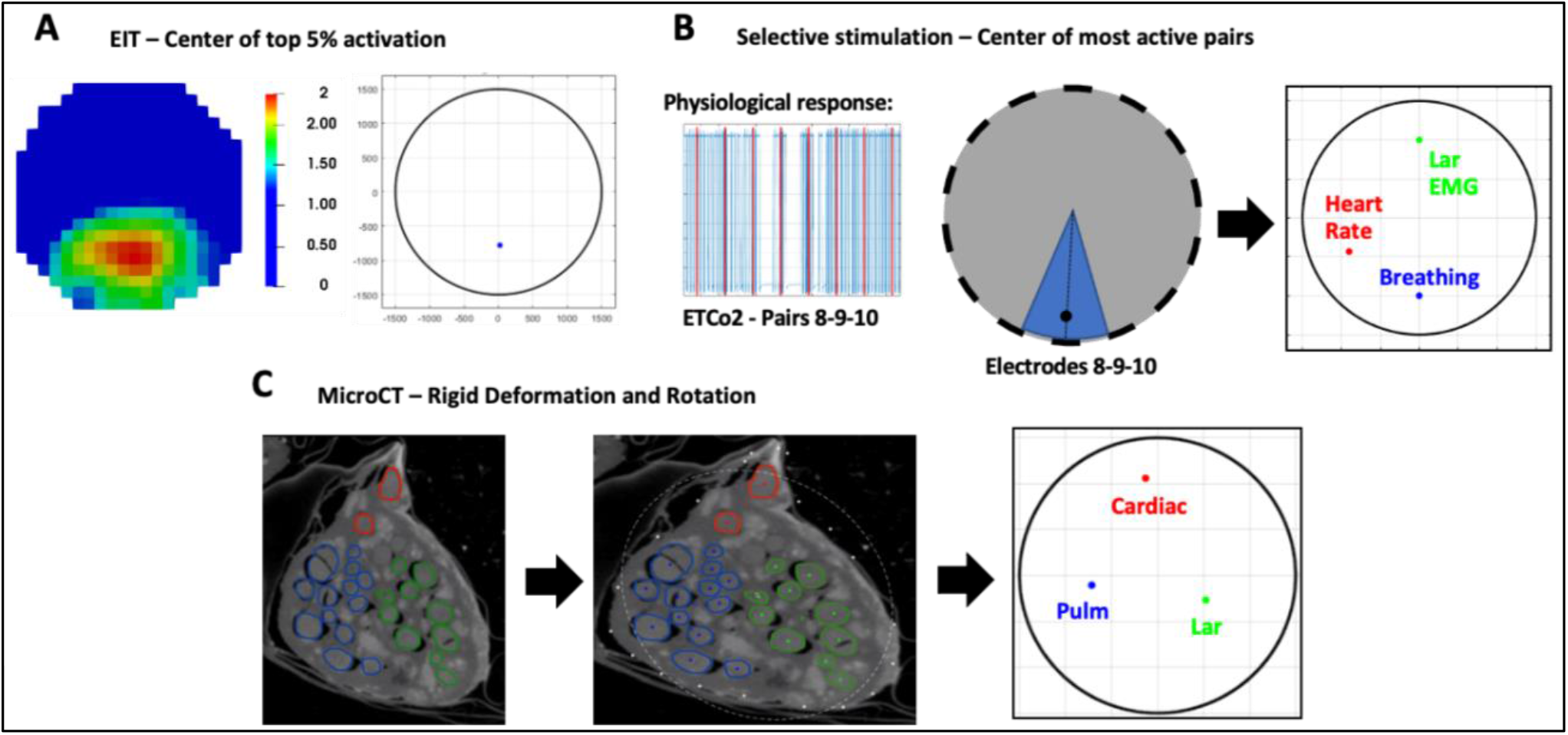
Image co-registration. **(A)** Voxels with top 5% magnitude from the reconstructed EIT image are selected to compute CoM of functional activation. The color scale is arbitrary units (Z score of relative change in the modulus of the impedance). **(B)** Electrode pair(s) inducing most localized and highest-intensity physiological response during selective stimulation were chosen to compute angular component of the center of mass for this technique; radial component is chosen as 2/3rds of nerve radius. **(C)** Centre of mass for each branch is computed from segmented MicroCT images which are then subject to rigid deformation and rotation for co-registration with the other methods.

MicroCT images underwent rigid deformation and rotation to have the same orientation and shape as EIT/SS data. The reconstruction of the SS and EIT images and CoMs were based on a generic circular cuff (Aristovich et al., 2018) so EIT images and sVNS locations appear on a perfectly circular geometry. Due to the fit of the cuff on the nerve and imperfect nerve geometry the vagus nerve as seen in microCT cross-sections is not perfectly circular. To allow for accurate comparison to the SS and EIT images and CoMs, the microCT cross-section had to be ‘deformed’ to fit the shape of a circle. The deformation of MicroCT images was performed as opposed to accurate modelling in SS and EIT to account for the fact that SS and EIT will be used in future for human studies, where accurate histology-based modelling is impossible; therefore, co-registration using the perfect circle would allow more realistic estimations of the accuracies relevant to the human nerve or chronic studies with SS and EIT. MicroCT images such as the one from Figure 4 were first rotated until the cuff opening was located at the same angle as in the EIT images and sVNS (on top at 90 degrees on the plane of the nerve cross-section), to match the coordinate systems; then, the microCT image was stretched until it formed a circular shape. Stretching as performed as rigid deformation, meaning that images were only elongated or shortened across the X/Y axis, to turn from ellipses into circles. This process was performed independently for each animal. Afterwards, CoM coordinates for each branch were computed by manually labelling the center locations of all fascicles related to a branch and then averaging individual center coordinates. The CoM error over all three branches was first compared in terms of cartesian distance and then split into radial and angular mismatch for easier interpretation. While EIT/SS techniques can be compared to their reference microCT data on individual nerves, grouping of CoM results over multiple nerves to analyze relative distribution of branches poses a challenge due to their different location in relation to the cuff angular orientation. A clusterization procedure (Ravagli et al., 2021) was adopted for EIT, SS and microCT to investigate the consistency of the distribution of the three functions (laryngeal, pulmonary and cardiac) over the cross-section of the nerve across animals. This procedure consisted in a rigid rotation of the CoM coordinates of all three organs until the dispersion of CoM from each branch among animals was minimized overall; being a rigid rotation, the relative position of each CoM within each animal remained constant. This procedure was implemented using a rotation matrix applied to the CoM coordinates at different angles until dispersion was minimized. Afterwards, the central coordinates of the laryngeal cluster were put on top to give some common reference to the panels of Figure 6. All quantities in this work were reported as mean ± standard deviation (SD) unless specified.

## 3 Results

### 3.1 Functional imaging of the fascicular organization of the vagus nerve *in vivo*

#### 3.1.1 EIT allows imaging of organ-specific activity within the vagus nerve

A SNR of 4.1±1.5, 3.1±0.16, and 3.1±0.65 with 120±32, 126±22, 126±22 δV traces (≈83, 87 and 87% of available) were used for image reconstruction for EIT of recurrent laryngeal, pulmonary, and cardiac activity, respectively (N=4). These revealed distinct regions corresponding to the three functions (Figure 5C). Locations of peak functional activity identified by FN-EIT matched post-mortem microCT tracing with an accuracy of <25% of nerve diameter, equally split between the radial and angular component.

**Figure 5:**
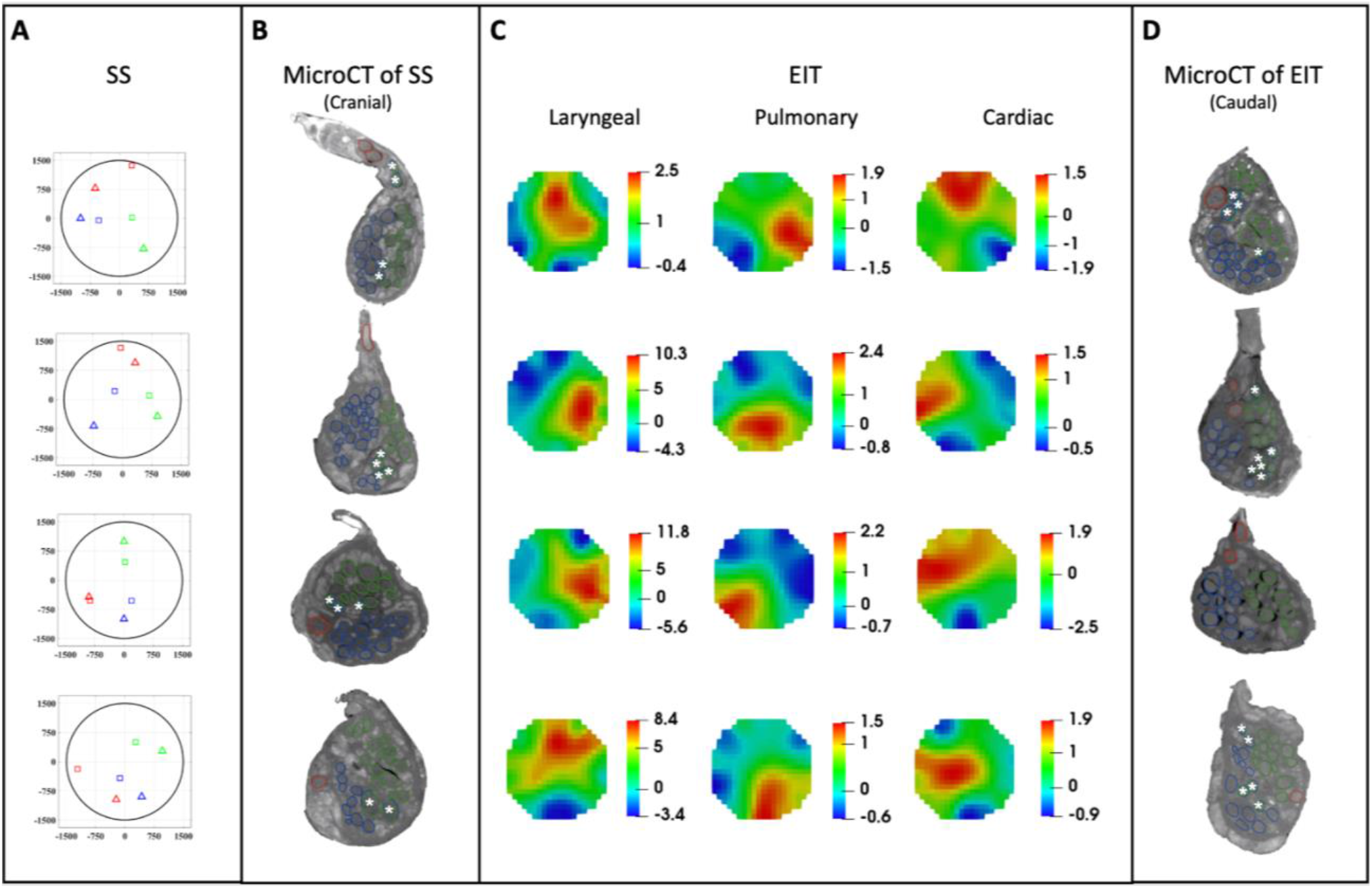
EIT, SS and microCT results (N=4) rotated with respect to the cuff opening (top) (laryngeal in green, pulmonary in blue, and cardiac in red). (**A**) Selective stimulation (SS) CoM plots. Triangles = SS, Squares = microCT. (**B**) MicroCT cross-sections of the center of SS cuff with segmented fascicles. Merged pulmonary and recurrent laryngeal fascicles are demarcated with a white asterisk. (**C**) EIT images for each fascicular group in the nerves. The color scale is arbitrary units (Z score of relative change in the modulus of the impedance). (**D**) MicroCT cross-sections of the center of EIT cuff with segmented fascicles. Merged pulmonary and recurrent laryngeal fascicles are demarcated with a white asterisk. **A-D** rotated with respect to the cuff opening at 12 o’clock.

#### 3.1.2 Selective stimulation allows for localization of organ-specific activity

Stimulation parameters which achieved spatial selectivity (physiological readout observed in less than 7 out of 14 pairs) were 0.4-0.8/1-2/0.1-0.2 mA, 50 μs/1 ms/50 μs pulse width, at 20 Hz for pulmonary, cardiac and laryngeal stimulation, respectively. This caused significant changes in the four nerves of −10.1±4.4%, −38.6±13.7%, and an increase of >10x baseline in the ECG, respiratory frequency, and EMG RMS value, respectively. These changes in peripheral activity were all statistically significant (p<0.05) and were elicited in 4±2 consecutive pairs out of 14 serial radial electrode pair sites (Figure 5A) in all animals.

### 3.2 MicroCT as a gold standard for imaging structural fascicular organization

MicroCT was used to image the vagus nerve from the cervical level to distal cardiac, recurrent laryngeal, and pulmonary branches. Lengths of up to 28cm could be imaged with an isotropic voxel size of 7 μm, digitally aligned between overlapping regions between scans, and segmented and traced with Neurolucida 360 (Version 2021.1.3, MBF Bioscience LLC, Williston, VT USA) (N=4, 24.25 cm ± 1.66 in length) (Figures 5B and D and 7). There was clear distinguishability of the fascicles from the rest of the nerve soft tissue (Thompson et al., 2020). Selective stimulation and FN-EIT were undertaken with different cuffs at one level each, 3 and 5.5 cm inferior to the nodose ganglion, respectively (termed “cranial” and “caudal” cuff level); microCT analysis was undertaken at both levels. All contoured and segmented regions identified as fascicles in a cross-section of the nerve from the middle of both cranial and caudal cuff placements were validated against the histology with H&E. No regions of the nerve other than fascicles were segmented. The number of fascicles was consistent between the microCT and respective histology images (Supplementary Figures 4 and 5). On average, the superior and inferior cardiac branches of the vagus nerve had an area of 0.16±0.1 mm^2^ and 0.12±0.04 mm^2^ and contained 1.25±0.5 and 1 fascicle(s) with an area of 0.024±0.01 mm^2^ and 0.023±0.01 mm^2^, respectively. The recurrent laryngeal branch had an area of 0.42±0.2 mm^2^ and contained 6.75±2.6 fascicles with an average area of 0.014±0.01 mm^2^. Pulmonary branches had an average area of 0.18±0.08 mm^2^ and contained 5±1 fascicles with an average area of 0.02±0.01 mm^2^. For full morphometric analysis of the vagus nerve and fascicles at the two cuff levels and of each branch, please see the supplementary files. On average, after fixation in formalin, the distance from the cervical level (taken as the top of the first, cranial cuff) to the branches of the vagus nerve were as follows: superior cardiac branch (8.25 cm ± 1.55), inferior cardiac branch (11.25 cm ± 2.96), recurrent laryngeal branch (20.63 cm ± 1.89), pulmonary branches (22.25 cm ± 2.10, 24.25 cm ± 1.66 and 26 cm). The length of the vagus nerve between the superior and inferior cardiac branches was 3.00 cm ± 1.58 and between the recurrent laryngeal branch and pulmonary branches was 2.00 cm ± 0.91 and 3.63 cm ± 0.63, respectively. Only one nerve had a third pulmonary branch with visible fascicles which was 26 cm from the cervical level and 5 cm from the recurrent laryngeal branch. During dissection, it appeared most nerves had additional pulmonary or cardiac branches; however, it was confirmed in the microCT images that some appearing to be branches during dissection of the nerve were merely connective tissue with no fascicles present. This may explain the so-called variation in anatomy and branching observed by eye during microdissection (Boyd, 1949; Hammer et al., 2018b; Settell et al., 2020; Tubbs et al.).

### 3.3 Consistent organization of cardiac, pulmonary, and recurrent laryngeal structural and functional fascicular groups across animals

Functional activity and anatomy for the three end-organs correlated largely exclusively to different regions of the cross-section of the left vagus nerve in the proximal neck. The functional and structural organization was consistent between animals (N=4) and across the three techniques. Overall, the cardiac, pulmonary and laryngeal fascicles were located ventromedially, dorsomedially, and ventrolaterally to laterally, respectively, for all three techniques and all four nerves at mid-cervical level (Figures 5A-D, 6, 7 and 8). There were 30.5±4.4 fascicles present per nerve. Of these, there were 1.4±0.5, 17.8±5.7 and 15.0±2.6 fascicles correlating to cardiac, pulmonary, and laryngeal activity, respectively. The cardiac correlating fascicles were exclusive (Supplementary Figure 6); 3.6±2.1 correlated to both pulmonary and recurrent laryngeal function (mean±SD). The point at which pulmonary and recurrent laryngeal fascicles began to merge varied between the four animals: the first merger between these two organ-specific groups took place at 4.5, 8, 9.5 and 16 cm from the nodose ganglion in the four nerves, respectively (9.5±4.8 cm). Both the recurrent laryngeal and pulmonary branches have not yet branched, and thus fascicles were present in the nerve, 25±1.5 cm from the nodose ganglion. The distance between the first merge between organ-specific fascicles and the point along the vagal trunk prior to branching of recurrent laryngeal and pulmonary branches was 15.75±4.8 cm. Angular separation with respect to the laryngeal fascicles was 118±37° and 114±77° for EIT, 125±42° and 128±36° for selective stimulation (SS), 102±28° for cranial cuff level microCT, and 146±22° and 92±27° for caudal cuff level microCT for pulmonary and cardiac, respectively (p<0.05 except 0.06 for cardiac laryngeal FN-EIT). After clusterization and CoM analysis, dispersion of fascicles around their center was 408 μm, 361 μm, 156 μm, and 182 μm for EIT, SS, cranial and caudal level microCT, respectively (Figure 6). The area of the cervical vagus nerve was 3.5±0.3 and 2.9± 0.40 mm^2^, for cranial and caudal cuffs, respectively. The areas of the regions containing the correlating fascicular groups was equal and greater for pulmonary and laryngeal and ≈10x smaller for cardiac region (32.4±5.8, 29.4±3.5 and 3.5±1.4% of total area, respectively). Overlap between pulmonary and laryngeal function occurred in 6.5±3.8% of the nerve area containing fascicles (62.9±10.1%). The datasets generated for this study can be found at Pennsieve Discover (https://discover.pennsieve.io/) with the title “Organotopic Organization of the Porcine Vagus Nerve – MicroCT, EIT and Selective Stimulation” (doi: 10.26275/hmwa-nqdu).

**Figure 6:**
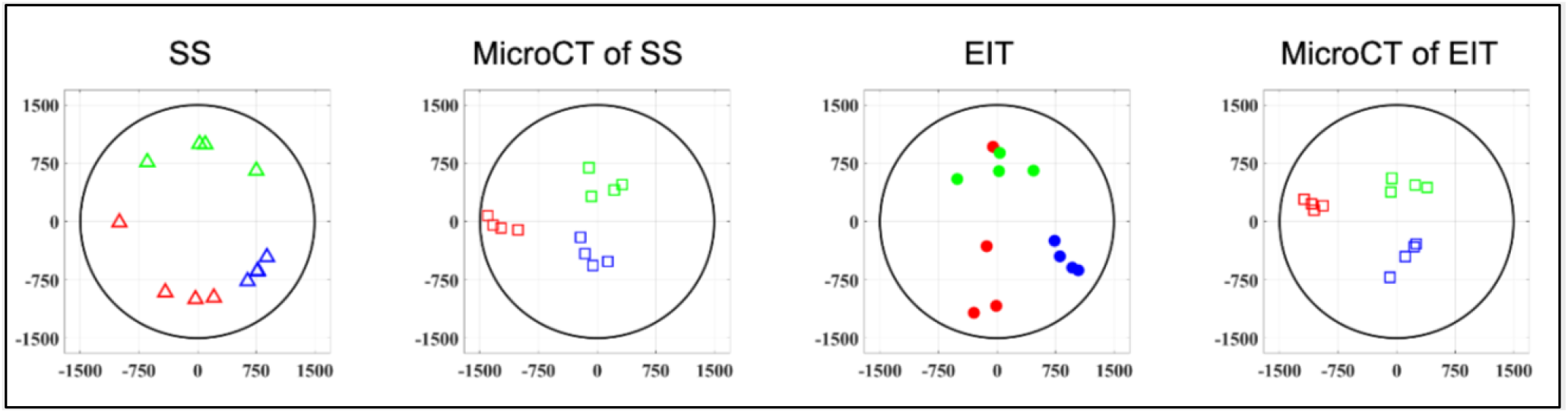
Clustered CoMs of the fascicular groups of each nerve for each technique rotated with respect to the laryngeal fascicles (N=4) (laryngeal in green, pulmonary in blue and cardiac in red).

**Figure 7:**
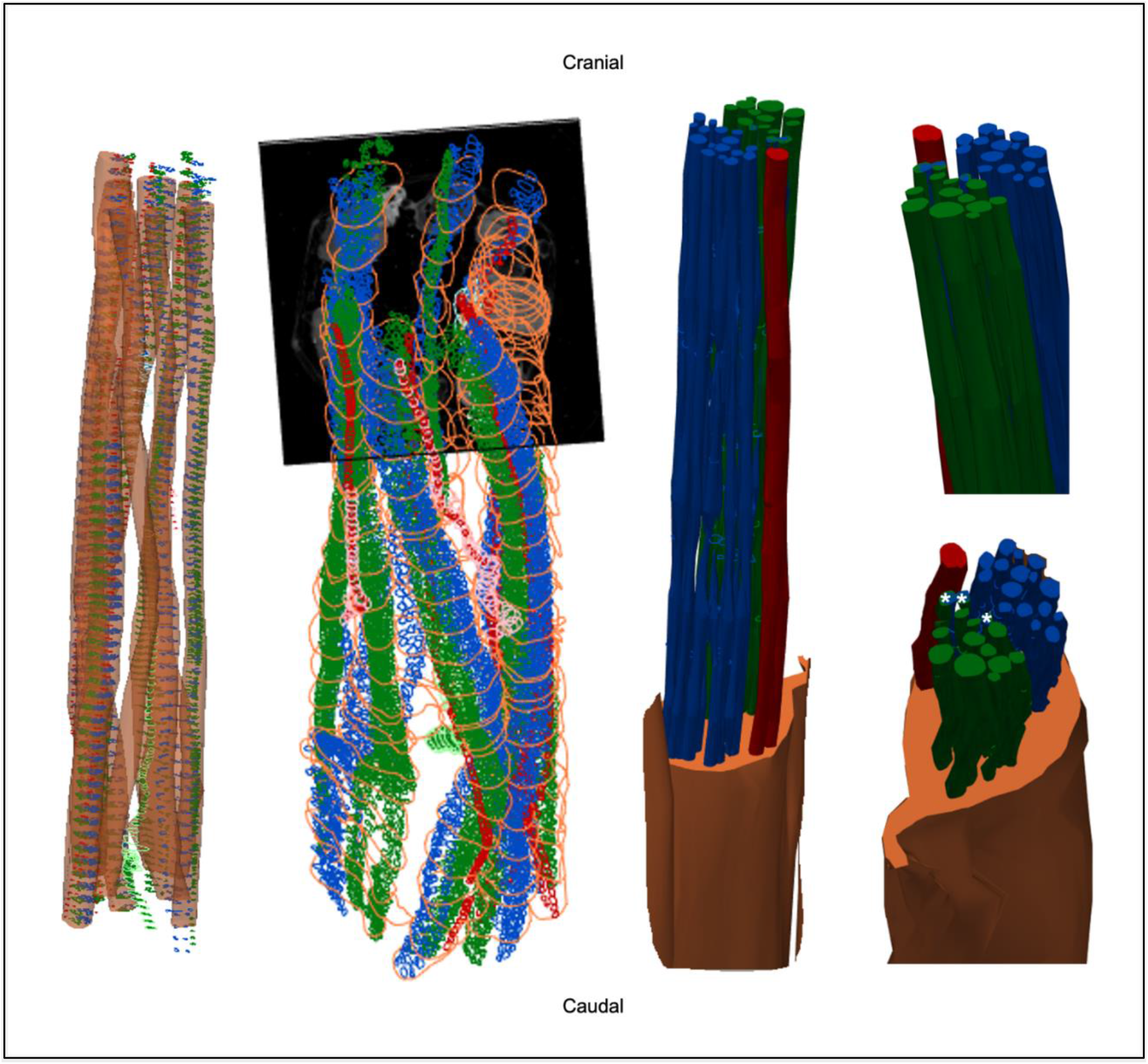
3D shelled segmentation of fascicles in the nerve. An example of the segmentation of the fascicles in the nerve within a stitched scan (all four scans stitched together) with contours placed every 50 to 100 sections throughout the length of the six segments of nerve. The contours can be joined together (right) by 3D shelling data of a certain label (i.e. cardiac or pulmonary or vagus nerve). Recurrent laryngeal is shown in green, pulmonary in blue, cardiac in red and the vagus trunk in orange. Merged fascicles between pulmonary and recurrent laryngeal are marked with an asterisk in the bottom right image.

**Figure 8:**
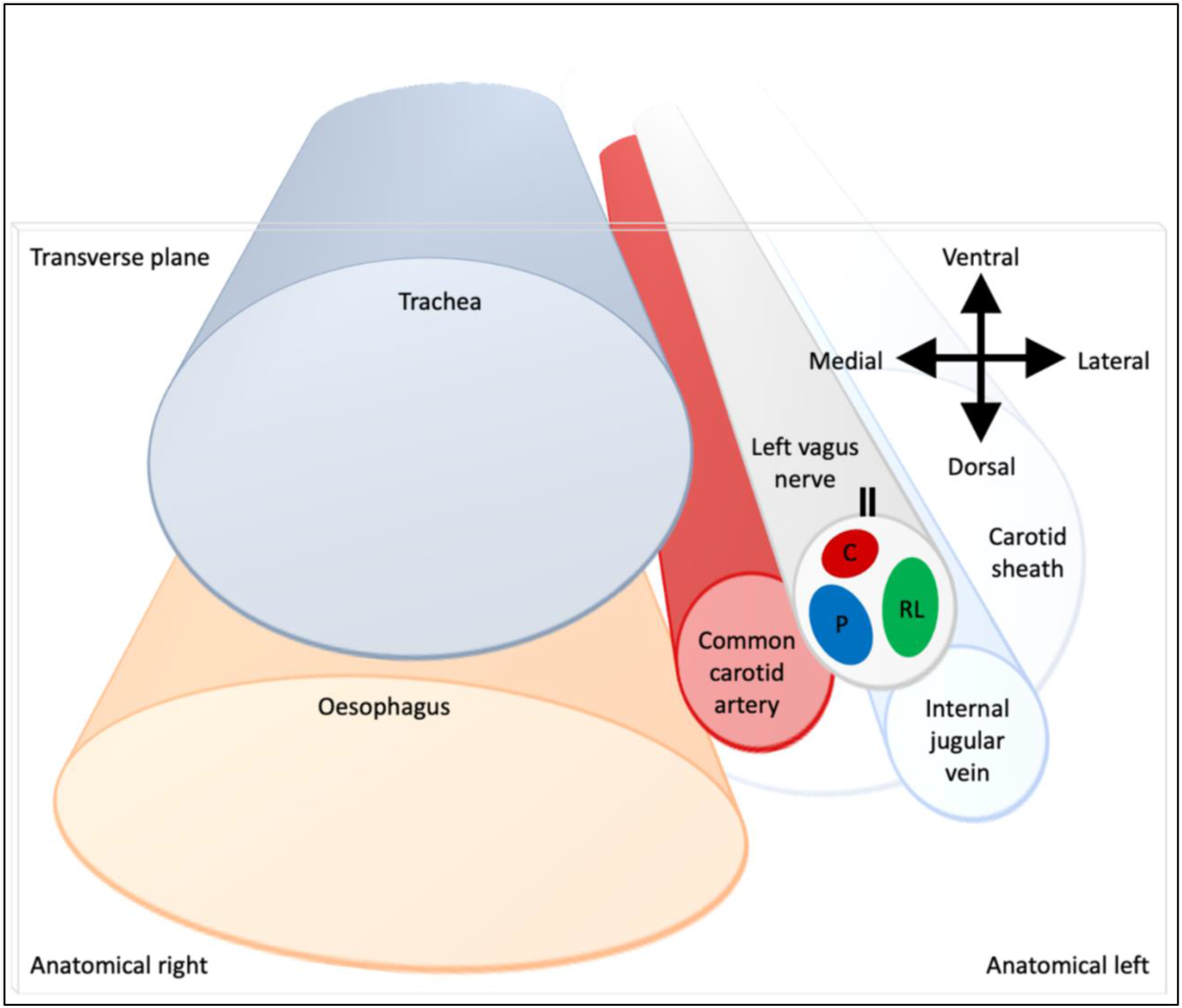
Schematic of anatomical location of fascicular regions of the left vagus nerve within the pig with a caudal to cranial view. The opening of the cuff is indicated with a double black line and the fascicular groups have a clockwise order of laryngeal (green), pulmonary (blue) and cardiac (red). V = ventral, VM = ventromedial, M = medial, DM = dorsomedial, D = dorsal, DL = dorsolateral, L = lateral, and VL = ventrolateral.

## 4 Discussion

The three methods have enabled the fascicular organization of the cervical vagus nerve at the level of VNS cuff placement to be deciphered for the first time. It was possible to identify three spatially separated fascicular groups which correlated with cardiac, pulmonary and laryngeal (thoracic) activity with FN-EIT and selective VNS, and this correlated closely with microCT tracing of the organ branches from their entry into the vagus nerve up to the cervical level (N=4, ~28 cm). The cervical vagus nerve is arranged organotopically with respect to these three fascicular groups. These findings were consistent between nerves and the functional and structural imaging techniques. In a cross-section, if recurrent laryngeal is placed at the top (12 o’clock), pulmonary and cardiac follow in clockwise order (c. 5 o’clock and 9 o’clock, respectively).

The vagus nerve innervates the heart, tracheobronchial tree and lungs and the muscles of the larynx in addition to the stomach, esophagus, pancreas, liver and gastrointestinal tract (Thompson et al., 2019). In this study, only thoracic fascicular groups were studied as EIT can only be undertaken by imaging differences over time. This requires averaging to a repeated trigger or electrical stimulation to an organ with fast myelinated fibers; the ECG, respiration or laryngeal electrically evoked activity appeared to give the most suitable opportunity. Correlation to subdiaphragmatic organs was not undertaken and is currently in progress. It seems probable that some of the fascicles within the three identified fascicular regions also contained subdiaphragmatic innervation as well; the three identified organs appeared to contribute to ~95% of observed fascicles. As subdiaphragmatic fibers are about one third of the cross-sectional area and number of fascicles of the cervical vagus nerve in pigs as well as humans (Pelot et al., 2020), it seems probable that subdiaphragmatic fibers may share some of the fascicles supporting thoracic function identified in this study.

On the microCT studies, it was possible to identify fascicles but not individual fibers. It appeared that some peripheral fascicles merged into others and then these combined fascicles diverged more proximally into two or more. If so, the more proximal divergent fascicles were considered to subserve the function of its parent fascicle and included in subsequent tracing of the fascicular group up the nerve. This was on the hypothesis that fibers pass between fascicles during this cross over. This did not pose any uncertainty for cardiac fascicles which remained distinct throughout. However, this may have led to some false positives or overreporting of proximal fascicles for the other fascicular groups, as the diverging fascicle need not necessarily have subserved the parent function. Without tracing of individual fiber function, this effect could not have been definitely characterized, and may have led to an overestimate of fascicles subserved by pulmonary or laryngeal function. In principle, this could have been addressed by fascicle tracing using neural tracers, but no reliable technique is described for the length of nerve characterized in this work in the pig. Only the left vagus nerve was used in this study to correspond with the predominant use of left vagi in clinical VNS applications: the left VNS is reported to cause fewer off-target cardiac effects, when treating epilepsy, including bradycardia or asystole These are hypothesized to be mediated primarily by the right vagus (Howland, 2014). In keeping with somatotopic organization of somatic nerves and organotopic organization of the left vagus nerve and that the left and right vagus provide innervation to the same thoracic end targets, it seems reasonable to suspect that there would be a similar organization of fascicles in the right vagus nerve.

These findings support the hypothesis based on somatic fascicle organization that cervical vagus nerves are organized organotopically. Fascicles are delimited by the perineurium. This is formed by cellular layers and collagen fibers which form a sleeve-like tubular sheath surrounding the fascicles (William K Ovalle and Nahirney, 2021). Its function is to provide tensile strength and elasticity to the nerve. It acts as a diffusion barrier to irritants and maintains homeostasis of the endoneurium and constant intrafascicular pressure (Peltonen et al., 2013; David J Magee and Manske, 2021). Its role in fiber organization is not entirely clear but rather seems to be a convenient envelope that allows for the intermingling or movement of fibers from one fascicle to another whilst inadvertently assisting in the maintenance of fiber organization. In somatic nerves, fascicles appear to maintain a somatotopic organization and may have a simpler parallel cable like or more intermingling plexiform arrangement (Sunderland, 1978; Jabaley et al., 1980). In general, there appears to be discrete somatotopic clustering of fibers distally toward the innervated muscles with intermingling of fibers and fascicles proximally (Langley and Hashimoto, 1917; McKinley, 1921; Sunderland, 1945). Tracing of fibers in somatic nerves indicated that, despite some movement of fibers between fascicles, they tended to remain clustered together within the proximal fascicle (Jabaley et al., 1980; Chow et al., 1985; Watchmaker et al., 1991). Within the ANS, there has been development of methods to enable imaging and subsequent tracing of the fascicular anatomy of the vagus nerve (Kolluru et al., 2021; Settell et al., 2021); however, until now, to our knowledge, no conclusive tracing of fascicles within the ANS has been performed. The arrangement in this study in the pig appears to be both cable-like and plexiform, but with the maintenance of discrete fascicular groupings of the three organs at the cervical level, in a fashion similar to that in somatic nerve but with respect to organs.

The anatomical knowledge of the fascicular organization in the cervical vagus nerve is an important novel scientific finding alone, but in addition, it could aid the following fields in neuroscience: 1) More precise investigations of healthy neural control of visceral organs as the neural efferent and afferent signals within the vagus nerve can be separated and organ-specific fibers identified; As an example, the cardiac neural control is of great interest, especially with respect to afferent VS efferent traffic and how this could be leveraged to understand and precisely control the autonomic system remodeling in chronic heart failure (Ardell et al., 2015, 2017; Ardell and Armour, 2016). 2) The investigation of neurological disease and dysfunction of the autonomic neural control can be greatly enhanced by studying fiber and cell function, structure and neurodegeneration overlayed on the organotopic functional map. This will be useful in studies of pain (Busch et al., 2013), interactions between sympathetic and parasympathetic systems (Deuchars et al., 2018; Bonaz et al., 2021; Kamiya et al., 2021), and mechanisms of various conditions caused by autonomic nervous system dysfunction. 3) Organ-specific invasive autonomic neurophysiology is now possible using electric (Fitchett et al., 2021), optical (Fontaine et al., 2021), or chemical (Ahmed et al., 2022) methods, by isolating the organ-specific fascicles from the vagus. In some cases, this could be the only method as it is impossible to localize and isolate all organ-specific fibers at the organ level due to sprouting (Jensen et al., 2013) or joining of other neural structures such as ganglia (Bratton et al., 2012). 4) The studies of ephaptic coupling (Anastassiou et al., 2011) and interactions would be greatly aided by the knowledge of the organ-specific fiber locations. As an example, the mechanisms of visceral pain could be studied with a clear prediction mechanism (Finnerup et al., 2021).

These results also lend hope to more clinical applications: the improvement of nerve repair and regeneration, microsurgery, and possible use of selective VNS in the future. The latter is currently accomplished with stimulation of the entire cervical vagus nerve; this indiscriminately modulates all organs supplied and consequent unwanted side-effects, such as cough, dyspnea, hoarseness, shortness of breath and bradycardia (Mulders et al., 2015), limit therapeutic efficacy (Howland, 2014; Ripplinger, 2017; Thompson et al., 2019). In principle, this could be avoided by spatially-selective stimulation of individual fascicles with knowledge of the fascicular organization of the vagus. This can allow expansion of VNS from its current use in the treatment of drug-resistant epilepsy and depression (Nemeroff et al., 2006; Fisher et al., 2021) to cardiovascular disorders and heart failure, lung injury, asthma, sepsis, arthritis, diabetes, pain management, and even immune function (Chakravarthy et al., 2015; Mehmed, 2015; Asad and Stavrakis, 2019; Drewes, 2021; Li et al., 2021; Marsal et al., 2021). The recurrent laryngeal fascicles identified at the cervical level accounted for roughly a half of fascicles present, correlating with previous studies (Settell et al., 2020). Avoidance of undesired stimulation of vagal outflow to the larynx alone could improve tolerability and efficacy of VNS.

This work was in the pig, a preclinical model for humans (Wolthuis et al., 2016). Pigs have a multifascicular vagus nerve and are the closest experimental animal to human vagus nerves in terms of anatomy, the diameter and length, the general morphology and its pathophysiology. The pig vagus nerve has been used in a number of studies for the analysis and investigation of neuromodulation parameters and effects, and the study of the anatomy and morphology of the nerve for eventual progression to humans (Rajendran et al., 2016; Wolthuis et al., 2016; Nicolai et al., 2020; Settell et al., 2020, 2021). *Ex vivo* microCT studies are in progress in humans in our group to ascertain if human vagal mapping is similar in conjunction with implantable human nerve cuff development. With this, relative mapping and alignment could be performed *in vivo* using physiological readouts (EMG, ECG, respiratory rate) to confirm the respective positions of the three regions upon performing spatially-selective VNS with the multi-electrode cuff. In addition, EIT can be used to image functional activity in the nerve corresponding to the organ of interest. This could enable selective stimulation in humans for which there is no current technique for *in vivo* fascicle or nerve activity visualization. It should be possible to identify mapping of the other organs innervated by the vagus nerve using microCT and anatomical tracing. Functional *in vivo* determination is more challenging for subdiaphragmatic organs which are supplied only with unmyelinated nerve and lack the clear phasic activity present for heart and lungs used in this study. It may nevertheless be possible to corroborate microCT mapping using phasic adaptive electrical stimulation (Tarotin et al., 2019) and selective stimulation in the cervical vagus nerve providing there is an end organ readout, such as venous noradrenaline in the splenic vein.

Conductivity variations reconstructed by EIT cover a large portion of the volume. This is a known effect of the mathematical regularization process, which creates a spatial filtering effect, and it is unavoidable as regularization is necessary to reconstruct the image by compensating for the mathematical problem being ill-posed and ill-conditioned. For circular geometries, the resolution can be roughly defined as 5-10% of the diameter, and it is usually possible to distinguish two objects located <10% of the diameter of the circle (Adler and Holder, 2021), therefore it is within the fascicular size for this study. However, the fascicular distinguishability was outside the scope of this study; the focus of this study in regard to nerve EIT was the localization of neural activity (single or multi-fascicular) within the nerve and we have found the accuracy of such localization, which was <25% nerve diameter (CoM co-localization error). This is within the specified requirements for sVNS as it matches the achievable selective stimulation accuracy (Aristovich et al., 2021). However, this can be further improved since part of the localization error may be due to the co-registration process with MicroCT images.

EIT is known to be a technique for which achieving a good enough SNR is challenging (Aristovich et al., 2018), which is the reason why coherent averaging is performed. Typically, because of the regularization needed for image reconstruction, it is possible to achieve acceptable image qualities when the impedance change is at least at the same level of noise (SNR>=1), but of course higher levels are preferred (e.g. SNR>=3). In this work, multiple technical solutions were employed to maximize SNR and detection of impedance changes, and to avoid the acquisition of artefacts. EIT was performed in time-difference mode, which greatly attenuates the residual effect of unbalanced contact impedance across electrodes, which remains static over the duration of action potential propagation (several milliseconds) (Chapman et al., 2019). EIT was performed at 6 kHZ frequency, which was identified as the optimal frequency for maximal SNR in EIT recordings in peripheral nerves (Aristovich et al., 2018). Other sources of electrical activity such as ECG, EMG, and brain waves could in theory influence EIT recordings; however, they are usually low-frequency (<1-2 kHz) and were filtered out by our bandpass filter. More so, i) most of the flow of current was confined to the inside of the EIT nerve cuff, ii) coherent averaging performed over time windows designated by the target branch would average out most confounders, and iii) for spontaneous recordings, RMS conversion of EIT signals would negate any possible low-frequency contribution around the EIT carrier frequency of slow mechanical confounders such as breathing. This work also benefits from prior knowledge and control studies (Aristovich et al., 2018) which showed the stability of EIT recordings to artefacts such as electrical-, EMG- and motion-related by analysis independence of percentage impedance changes and by applying lidocaine to the nerve. We believe that the combination of the above controls and considerations eliminates any possibility of acquiring artefacts.

### 4.1 Conclusions

The left cervical vagus nerves of pigs were reproducibly organized with respect to cardiac, pulmonary, and recurrent laryngeal function. This supports the hypothesis that fascicles in the autonomic nervous system are organized, at least to a substantial extent, in an organotopic fashion. This supports the possibility of changing current practice in VNS with selective stimulation and increasing therapeutic efficiency by avoidance of off-target effects. The novel techniques of FN-EIT and trial-and-error selective stimulation show promise for *in vivo* imaging of functional fascicular organization and localization clinically in humans with an accuracy sufficient for targeted VNS.

## Author Contributions

Conceptualization: KA, DH

Methodology: NT, ER, SM

Investigation: NT, ER, SM, JP, KA

Visualization: NT, ER

Funding acquisition: KA, DH

Project administration: NT, ER, SM, KA, DH

Supervision: KA, DH

Writing – original draft: NT, ER, SM

Writing – review & editing: NT, ER, SM, JP, KA, DH

## Acknowledgments

Thank you to Maci Heal and Shane Baldwin, MBF Bioscience, for their assistance with Neurolucida 360 software, tools, and training. Thank you to David Goodwin, Royal Veterinary College, for his assistance with histology.

## Data Availability Statement

The datasets generated for this study can be found at Pennsieve Discover (https://discover.pennsieve.io/datasets/287) with the title “Organotopic Organization of the Porcine Vagus Nerve – MicroCT, EIT and Selective Stimulation” (doi: 10.26275/hmwa-nqdu).

## References

Adler, A., and Holder, D. (2021). Electrical Impedance Tomography: Methods, History and Applications. CRC Press.

Ahmed, U., Chang, Y.-C., Zafeiropoulos, S., Nassrallah, Z., Miller, L., and Zanos, S. (2022). Strategies for precision vagus neuromodulation. Bioelectronic Medicine 8, 9. doi: 10.1186/s42234-022-00091-1.

Anastassiou, C. A., Perin, R., Markram, H., and Koch, C. (2011). Ephaptic coupling of cortical neurons. Nat Neurosci 14, 217–223. doi: 10.1038/nn.2727.

Ardell, J. L., and Armour, J. A. (2016). Neurocardiology: Structure-Based Function. Compr Physiol 6, 1635–1653. doi: 10.1002/cphy.c150046.

Ardell, J. L., Nier, H., Hammer, M., Southerland, E. M., Ardell, C. L., Beaumont, E., et al. (2017). Defining the neural fulcrum for chronic vagus nerve stimulation: implications for integrated cardiac control. J Physiol 595, 6887–6903. doi: 10.1113/JP274678.

Ardell, J. L., Rajendran, P. S., Nier, H. A., KenKnight, B. H., and Armour, J. A. (2015). Central-peripheral neural network interactions evoked by vagus nerve stimulation: functional consequences on control of cardiac function. American Journal of Physiology-Heart and Circulatory Physiology 309, H1740–H1752. doi: 10.1152/ajpheart.00557.2015.

Aristovich, K., Donegá, M., Blochet, C., Avery, J., Hannan, S., Chew, D. J., et al. (2018). Imaging fast neural traffic at fascicular level with electrical impedance tomography: proof of principle in rat sciatic nerve. Journal of Neural Engineering 15, 056025. doi: 10.1088/1741-2552/aad78e.

Aristovich, K., Donega, M., Fjordbakk, C., Tarotin, I., Chapman, C. A. R., Viscasillas, J., et al. (2021). Model-based geometrical optimisation and in vivo validation of a spatially selective multielectrode cuff array for vagus nerve neuromodulation. Journal of Neuroscience Methods 352, 109079. doi: 10.1016/j.jneumeth.2021.109079.

Asad, Z. U., and Stavrakis, S. (2019). Vagus nerve stimulation for the treatment of heart failure. Bioelectronics in Medicine 2, 43–54. doi: 10.2217/bem-2019-0012.

Avery, J., Dowrick, T., Faulkner, M., Goren, N., Holder, D., Avery, J., et al. (2017). A Versatile and Reproducible Multi-Frequency Electrical Impedance Tomography System. Sensors 17, 280. doi: 10.3390/s17020280.

Bai, L., Mesgarzadeh, S., Ramesh, K. S., Huey, E. L., Liu, Y., Gray, L. A., et al. (2019). Genetic Identification of Vagal Sensory Neurons That Control Feeding. Cell 179, 1129–1143.e23. doi: 10.1016/j.cell.2019.10.031.

Beekman, R., and Visser, L. H. (2004). High-resolution sonography of the peripheral nervous system - a review of the literature. European Journal of Neurology 11, 305–314.

Bokil, H., Laaris, N., Blinder, K., Ennis, M., and Keller, A. (2001). Ephaptic Interactions in the Mammalian Olfactory System. J. Neurosci. 21, RC173–RC173. doi: 10.1523/JNEUROSCI.21-20-j0004.2001.

Bonaz, B., Sinniger, V., and Pellissier, S. (2021). Therapeutic Potential of Vagus Nerve Stimulation for Inflammatory Bowel Diseases. Frontiers in Neuroscience 15. Available at: https://www.frontiersin.org/articles/10.3389/fnins.2021.650971 [Accessed December 11, 2022].

Boyd, A. K. (1949). Vagotomy and the anatomic variations in the vagus nerve. The American Journal of Surgery 78, 4–14. doi: 10.1016/0002-9610(49)90178-6.

Bratton, B. O., Martelli, D., McKinley, M. J., Trevaks, D., Anderson, C. R., and McAllen, R. M. (2012). Neural regulation of inflammation: no neural connection from the vagus to splenic sympathetic neurons. Exp Physiol 97, 1180–1185. doi: 10.1113/expphysiol.2011.061531.

Busch, V., Zeman, F., Heckel, A., Menne, F., Ellrich, J., and Eichhammer, P. (2013). The effect of transcutaneous vagus nerve stimulation on pain perception – An experimental study. Brain Stimulation 6, 202–209. doi: 10.1016/J.BRS.2012.04.006.

Capllonch-Juan, M., and Sepulveda, F. (2020). Modelling the effects of ephaptic coupling on selectivity and response patterns during artificial stimulation of peripheral nerves. PLOS Computational Biology 16, e1007826. doi: 10.1371/journal.pcbi.1007826.

Carolus, A. E., Lenz, M., Hofmann, M., Welp, H., Schmieder, K., and Brenke, C. (2019). High-resolution in vivo imaging of peripheral nerves using optical coherence tomography: a feasibility study. Journal of Neurosurgery 132, 1907–1913. doi: 10.3171/2019.2.JNS183542.

Cartwright, M. S., Baute, V., Caress, J. B., and Walker, F. O. (2017). Ultrahigh-frequency ultrasound of fascicles in the median nerve at the wrist. Muscle Nerve 56, 819–822. doi: 10.1002/mus.25617.

Chakravarthy, K., Chaudhry, H., Williams, K., and Christo, P. J. (2015). Review of the Uses of Vagal Nerve Stimulation in Chronic Pain Management. Curr Pain Headache Rep 19, 54. doi: 10.1007/s11916-015-0528-6.

Chapman, C. A. R., Aristovich, K., Donega, M., Fjordbakk, C. T., Stathopoulou, T.-R., Viscasillas, J., et al. (2019). Electrode fabrication and interface optimization for imaging of evoked peripheral nervous system activity with electrical impedance tomography (EIT). Journal of Neural Engineering 16, 016001. doi: 10.1088/1741-2552/aae868.

Chow, J. A., Van Beek, A. L., Meyer, D. L., and Johnson, M. C. (1985). Surgical significance of the motor fascicular group of the ulnar nerve in the forearm. J Hand Surg Am 10, 867–872. doi: 10.1016/s0363-5023(85)80164-7.

David J Magee, and Manske, R. C. (2021). “Chapter 1 Principles and Concepts,” in Orthopedic Physical Assessment (Elsevier), 1–72.e2. Available at: https://www.clinicalkey.com/#!/content/book/3-s2.0-B9780323522991000012.

De Ferrari, G. M., Stolen, C., Tuinenburg, A. E., Wright, D. J., Brugada, J., Butter, C., et al. (2017). Long-term vagal stimulation for heart failure: Eighteen month results from the NEural Cardiac TherApy foR Heart Failure (NECTAR-HF) trial. International Journal of Cardiology 244, 229–234. doi: 10.1016/j.ijcard.2017.06.036.

Deuchars, S. A., Lall, V. K., Clancy, J., Mahadi, M., Murray, A., Peers, L., et al. (2018). Mechanisms underpinning sympathetic nervous activity and its modulation using transcutaneous vagus nerve stimulation. Experimental Physiology 103, 326–331. doi: 10.1113/EP086433.

Dowrick, T., Sato Dos Santos, G., Vongerichten, A., and Holder, D. (2015). Parallel, multi frequency EIT measurement, suitable for recording impedance changes during epilepsy. Journal of Electrical Bioimpedance 6, 37–43. doi: 10.5617/JEB.2573.

Drewes, A. M. (2021). Treatment of Complications to Diabetic Autonomic Neuropathy With Vagus Nerve Stimulation. clinicaltrials.gov Available at: https://clinicaltrials.gov/ct2/show/NCT04143269 [Accessed October 12, 2021].

Finnerup, N. B., Kuner, R., and Jensen, T. S. (2021). Neuropathic Pain: From Mechanisms to Treatment. Physiological Reviews 101, 259–301. doi: 10.1152/physrev.00045.2019.

Fisher, B., DesMarteau, J. A., Koontz, E. H., Wilks, S. J., and Melamed, S. E. (2021). Responsive Vagus Nerve Stimulation for Drug Resistant Epilepsy: A Review of New Features and Practical Guidance for Advanced Practice Providers. Frontiers in Neurology 11, 1863. doi: 10.3389/fneur.2020.610379.

Fitchett, A., Mastitskaya, S., and Aristovich, K. (2021). Selective Neuromodulation of the Vagus Nerve. Frontiers in Neuroscience 15, 600. doi: 10.3389/FNINS.2021.685872/BIBTEX.

Fontaine, A. K., Futia, G. L., Rajendran, P. S., Littich, S. F., Mizoguchi, N., Shivkumar, K., et al. (2021). Optical vagus nerve modulation of heart and respiration via heart-injected retrograde AAV. Sci Rep 11, 3664. doi: 10.1038/s41598-021-83280-3.

Hammer, N., Löffler, S., Cakmak, Y. O., Ondruschka, B., Planitzer, U., Schultz, M., et al. (2018a). Cervical vagus nerve morphometry and vascularity in the context of nerve stimulation - A cadaveric study. Scientific reports 8, 7997. doi: 10.1038/s41598-018-26135-8.

Hammer, N., Löffler, S., Cakmak, Y. O., Ondruschka, B., Planitzer, U., Schultz, M., et al. (2018b). Cervical vagus nerve morphometry and vascularity in the context of nerve stimulation - A cadaveric study. Sci Rep 8. doi: 10.1038/s41598-018-26135-8.

Hope, J., Braeuer, B., Amirapu, S., McDaid, A., and Vanholsbeeck, F. (2018). Extracting morphometric information from rat sciatic nerve using optical coherence tomography. JBO 23, 116001. doi: 10.1117/1.JBO.23.11.116001.

Howland, R. H. (2014). Vagus Nerve Stimulation. Curr Behav Neurosci Rep 1, 64–73. doi: 10.1007/s40473-014-0010-5.

Isabella, A. J., Stonick, J. A., Dubrulle, J., and Moens, C. B. (2021). Intrinsic positional memory guides target-specific axon regeneration in the zebrafish vagus nerve. Development 148, dev199706. doi: 10.1242/dev.199706.

Jabaley, M. E., Wallace, W. H., and Heckler, F. R. (1980). Internal topography of major nerves of the forearm and hand: A current view. The Journal of Hand Surgery 5, 1–18. doi: 10.1016/S0363-5023(80)80035-9.

Jehl, M., Dedner, A., Betcke, T., Aristovich, K., Klofkorn, R., and Holder, D. (2015). A Fast Parallel Solver for the Forward Problem in Electrical Impedance Tomography. IEEE Transactions on Biomedical Engineering 62, 126–137. doi: 10.1109/TBME.2014.2342280.

Jensen, K. J., Alpini, G., and Glaser, S. (2013). Hepatic Nervous System and Neurobiology of the Liver. Compr Physiol 3, 655–665. doi: 10.1002/cphy.c120018.

Kamiya, A., Hiyama, T., Fujimura, A., and Yoshikawa, S. (2021). Sympathetic and parasympathetic innervation in cancer: therapeutic implications. Clin Auton Res 31, 165–178. doi: 10.1007/s10286-020-00724-y.

Kolluru, C., Subramaniam, A., Liu, Y., Upadhye, A., Khela, M., Druschel, L., et al. (2021). 3D imaging of the vagus nerve fascicular anatomy with cryo-imaging and UV excitation. in Three-Dimensional and Multidimensional Microscopy: Image Acquisition and Processing XXVIII (International Society for Optics and Photonics), 1164910. doi: 10.1117/12.2577037.

Langley, J. N., and Hashimoto, M. (1917). On the suture of separate nerve bundles in a nerve trunk and on internal nerve plexuses. J Physiol 51, 318–346. doi: 10.1113/jphysiol.1917.sp001805.

Li, S., Qi, D., Li, J., Deng, X., and Wang, D. (2021). Vagus nerve stimulation enhances the cholinergic anti-inflammatory pathway to reduce lung injury in acute respiratory distress syndrome via STAT3. Cell Death Discov. 7, 1–9. doi: 10.1038/s41420-021-00431-1.

Marsal, S., Corominas, H., Agustín, J. J. de, Pérez-García, C., López-Lasanta, M., Borrell, H., et al. (2021). Non-invasive vagus nerve stimulation for rheumatoid arthritis: a proof-of-concept study. The Lancet Rheumatology 3, e262–e269. doi: 10.1016/S2665-9913(20)30425-2.

Mastitskaya, S., Thompson, N., and Holder, D. (2021). Selective Vagus Nerve Stimulation as a Therapeutic Approach for the Treatment of ARDS: A Rationale for Neuro-Immunomodulation in COVID-19 Disease. Front. Neurosci. 15. doi: 10.3389/fnins.2021.667036.

McKinley, J. C. (1921). THE INTRANEURAL PLEXUS OF FASCICULI AND FIBERS IN THE SCIATIC NERVE. Archives of Neurology & Psychiatry 6, 377–399. doi: 10.1001/archneurpsyc.1921.02190040020002.

Mehmed, S. E. (2015). Effect of vagal stimulation in acute asthma. Clin Transl Allergy 5, P13. doi: 10.1186/2045-7022-5-S2-P13.

Mulders, D. M., de Vos, C. C., Vosman, I., and van Putten, M. J. A. M. (2015). The effect of vagus nerve stimulation on cardiorespiratory parameters during rest and exercise. Seizure 33, 24–28. doi: 10.1016/j.seizure.2015.10.004.

Nemeroff, C. B., Mayberg, H. S., Krahl, S. E., McNamara, J., Frazer, A., Henry, T. R., et al. (2006). VNS Therapy in Treatment-Resistant Depression: Clinical Evidence and Putative Neurobiological Mechanisms. Neuropsychopharmacol 31, 1345–1355. doi: 10.1038/sj.npp.1301082.

Nicolai, E. N., Settell, M. L., Knudsen, B. E., McConico, A. L., Gosink, B. A., Trevathan, J. K., et al. (2020). Sources of off-target effects of vagus nerve stimulation using the helical clinical lead in domestic pigs. J. Neural Eng. 17, 046017. doi: 10.1088/1741-2552/ab9db8.

Pelot, N. A., Goldhagen, G. B., Cariello, J. E., Musselman, E. D., Clissold, K. A., Ezzell, J. A., et al. (2020). Quantified Morphology of the Cervical and Subdiaphragmatic Vagus Nerves of Human, Pig, and Rat. Frontiers in Neuroscience 14, 1148. doi: 10.3389/fnins.2020.601479.

Peltonen, S., Alanne, M., and Peltonen, J. (2013). Barriers of the peripheral nerve. Tissue Barriers 1, e24956. doi: 10.4161/tisb.24956.

Plachta, D. T. T., Gierthmuehlen, M., Cota, O., Espinosa, N., Boeser, F., Herrera, T. C., et al. (2014). Blood pressure control with selective vagal nerve stimulation and minimal side effects. Journal of Neural Engineering 11, 036011. doi: 10.1088/1741-2560/11/3/036011.

Rajendran, P. S., Challis, R. C., Fowlkes, C. C., Hanna, P., Tompkins, J. D., Jordan, M. C., et al. (2019). Identification of peripheral neural circuits that regulate heart rate using optogenetic and viral vector strategies. Nat Commun 10. doi: 10.1038/s41467-019-09770-1.

Rajendran, P. S., Nakamura, K., Ajijola, O. A., Vaseghi, M., Armour, J. A., Ardell, J. L., et al. (2016). Myocardial infarction induces structural and functional remodelling of the intrinsic cardiac nervous system. The Journal of Physiology 594, 321–341. doi: 10.1113/JP271165.

Rangavajla, G., Mokarram, N., Masoodzadehgan, N., Pai, S. B., and Bellamkonda, R. V. (2014). Noninvasive Imaging of Peripheral Nerves. Cells Tissues Organs 200, 69–77. doi: 10.1159/000369451.

Raphael, D. T., Yang, C., Tresser, N., Wu, J., Zhang, Y., and Rever, L. (2007). Images of Spinal Nerves and Adjacent Structures With Optical Coherence Tomography: Preliminary Animal Studies. The Journal of Pain 8, 767–773. doi: 10.1016/j.jpain.2007.04.006.

Ravagli, E., Mastitskaya, S., Thompson, N., Aristovich, K. Y., and Holder, D. S. (2019). Optimization of the electrode drive pattern for imaging fascicular compound action potentials in peripheral nerve with fast neural electrical impedance tomography (EIT). Physiol. Meas. doi: 10.1088/1361-6579/ab54eb.

Ravagli, E., Mastitskaya, S., Thompson, N., Iacoviello, F., Shearing, P. R., Perkins, J., et al. (2020). Imaging fascicular organization of rat sciatic nerves with fast neural electrical impedance tomography. Nature Communications 11, 6241. doi: 10.1038/s41467-020-20127-x.

Ravagli, E., Mastitskaya, S., Thompson, N., Welle, E. J., Chestek, C. A., Aristovich, K., et al. (2021). Fascicle localisation within peripheral nerves through evoked activity recordings: A comparison between electrical impedance tomography and multi-electrode arrays. Journal of Neuroscience Methods 358, 109140. doi: 10.1016/j.jneumeth.2021.109140.

Rea, P. (2014). “Chapter 10 - Vagus Nerve,” in Clinical Anatomy of the Cranial Nerves, ed. P. Rea (San Diego: Academic Press), 105–116. doi: 10.1016/B978-0-12-800898-0.00010-5.

Ripplinger, C. M. (2017). From drugs to devices and back again: chemical vagal nerve stimulation for the treatment of heart failure. Cardiovasc Res 113, 1270–1272. doi: 10.1093/cvr/cvx142.

Schindelin, J., Arganda-Carreras, I., Frise, E., Kaynig, V., Longair, M., Pietzsch, T., et al. (2012). Fiji: an open-source platform for biological-image analysis. Nature Methods 9, 676–682. doi: 10.1038/nmeth.2019.

Settell, M. L., Kasole, M., Skubal, A. C., Knudsen, B. E., Nicolai, E. N., Huang, C., et al. (2021). In vivo visualization of pig vagus nerve ‘vagotopy’ using ultrasound. bioRxiv, 2020.12.24.424256. doi: 10.1101/2020.12.24.424256.

Settell, M. L., Pelot, N. A., Knudsen, B. E., Dingle, A. M., McConico, A. L., Nicolai, E. N., et al. (2020). Functional vagotopy in the cervical vagus nerve of the domestic pig: implications for the study of vagus nerve stimulation. J. Neural Eng. 17, 026022. doi: 10.1088/1741-2552/ab7ad4.

Sheehan, D. C., and Hrapchak, B. B. (1987). Theory and practice of histotechnology. Columbus, Ohio: Battelle Press.

Sheheitli, H., and Jirsa, V. K. (2020). A mathematical model of ephaptic interactions in neuronal fiber pathways: Could there be more than transmission along the tracts? Netw Neurosci 4, 595–610. doi: 10.1162/netn_a_00134.

Sunderland, S. (1945). The intraneural topography of the radial, median and ulnar nerves. Brain 68, 243–299.

Sunderland, S. (1978). Nerves and nerve injuries. 2nd ed. Edinburgh: Edinburgh: Churchill Livingstone.

Tarotin, I., Aristovich, K., and Holder, D. (2019). Model of Impedance Changes in Unmyelinated Nerve Fibers. IEEE transactions on bio-medical engineering 66, 471–484. doi: 10.1109/TBME.2018.2849220.

Thompson, N., Mastitskaya, S., and Holder, D. (2019). Avoiding off-target effects in electrical stimulation of the cervical vagus nerve: Neuroanatomical tracing techniques to study fascicular anatomy of the vagus nerve. Journal of Neuroscience Methods 325, 108325. doi: 10.1016/j.jneumeth.2019.108325.

Thompson, N., Ravagli, E., Mastitskaya, S., Iacoviello, F., Aristovich, K., Perkins, J., et al. (2020). MicroCT optimisation for imaging fascicular anatomy in peripheral nerves. Journal of Neuroscience Methods 338, 108652. doi: 10.1016/J.JNEUMETH.2020.108652.

Tubbs, R. S., Loukas, M., Shoja, M. M., Blevins, D., Humphrey, R., Chua, G. D., et al. An unreported variation of the cervical vagus nerve: anatomical and histological observations. 3.

Vasudevan, S., Vo, J., Shafer, B., Nam, A. S., Vakoc, B. J., and Hammer, D. X. (2019). Toward optical coherence tomography angiography-based biomarkers to assess the safety of peripheral nerve electrostimulation. J. Neural Eng. 16, 036024. doi: 10.1088/1741-2552/ab1405.

Verlinden, T. J. M., Rijkers, K., Hoogland, G., and Herrler, A. (2016). Morphology of the human cervical vagus nerve: Implications for vagus nerve stimulation treatment. Acta Neurologica Scandinavica 133, 173–182. doi: 10.1111/ane.12462.

Watchmaker, G. P., Gumucio, C. A., Crandall, R. E., Vannier, M. A., and Weeks, P. M. (1991). Fascicular topography of the median nerve: a computer based study to identify branching patterns. J Hand Surg Am 16, 53–59. doi: 10.1016/s0363-5023(10)80013-9.

William K Ovalle, and Nahirney, P. C. (2021). “Chapter 5 Nervous Tissue,” in Netter’s Essential Histology (Elsevier), 109–139.

Wolthuis, A. M., Stakenborg, N., D’Hoore, A., and Boeckxstaens, G. E. (2016). The pig as preclinical model for laparoscopic vagus nerve stimulation. Int J Colorectal Dis 31, 211–215. doi: 10.1007/s00384-015-2435-z.

