## Supplementary Information for "Organotopic Organization of the Cervical Vagus Nerve"

Supplementary Materials

### 1 Supplementary Data

#### 1.1 MicroCT imaging of vagus nerves

##### 1.1.1 Matching neighboring nerve segments

The nerve was cut into 4 cm segments subsequent to the placement of sutures as landmarks across the cut region. The number of suture landmarks differs between the cut regions to ensure the correct neighboring segments were matched up (an additional measure to the nerve segments being arranged in the correct order). The last whole cross section visible in the last scan of a segment of nerve was identified and visualized with the first whole cross section visible in the first scan of the next segment of nerve. The suture landmarks were identified and the cross sections' orientation aligned to allow for further correlation of fascicles across the cut region. The size, number, position and pattern of fascicles were matched up between the two cross-sections of each cut region between nerve segments (Supplementary Figure 3). Additional anatomical landmarks, such as blood vessels, connective tissue or fat cells were also used to assist in ensuring correct correlation between cross sections. Occasionally, fascicles were in the process of merging or splitting across the cut region (three examples are shown in the cross sections of Supplementary Figure 3); however, with process of elimination with the other identified and matched fascicles, and viewing the scans till the respective ends where the whole nerve may not be visible, but the slight movement of fascicles of interest is, it can be deduced that these fascicles follow on from one another. The identified merged fascicle or split fascicles will be incorporated in the continuation of segmentation.

##### 1.1.2 Histology for validation

All contoured and segmented regions deemed to be fascicles in a cross-section of the nerve from the level of VNS cuff placement were validated against histology and were confirmed to have been correctly allocated and identified as being fascicular tissue within the nerve. No regions of the nerve other than fascicles were segmented. The number of fascicles was consistent between the microCT and respective histology images (Supplementary Figures 4 and 5).

### 2 Supplementary Figures and Tables

#### 2.1 Supplementary Figures

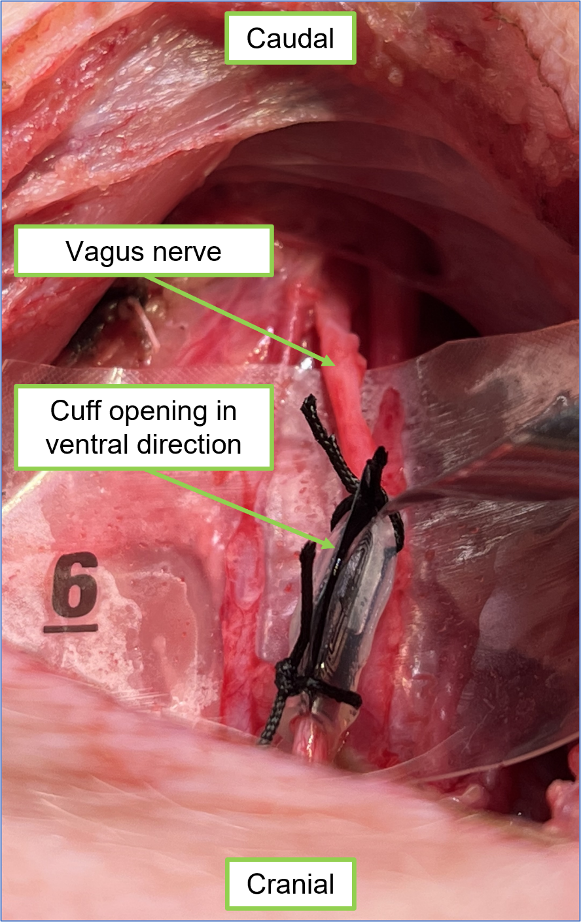

**Supplementary Figure 1. Placement of cuff with opening in ventral direction.** The cuff was placed on the nerve with the opening facing the ventral direction – toward the surgical opening.

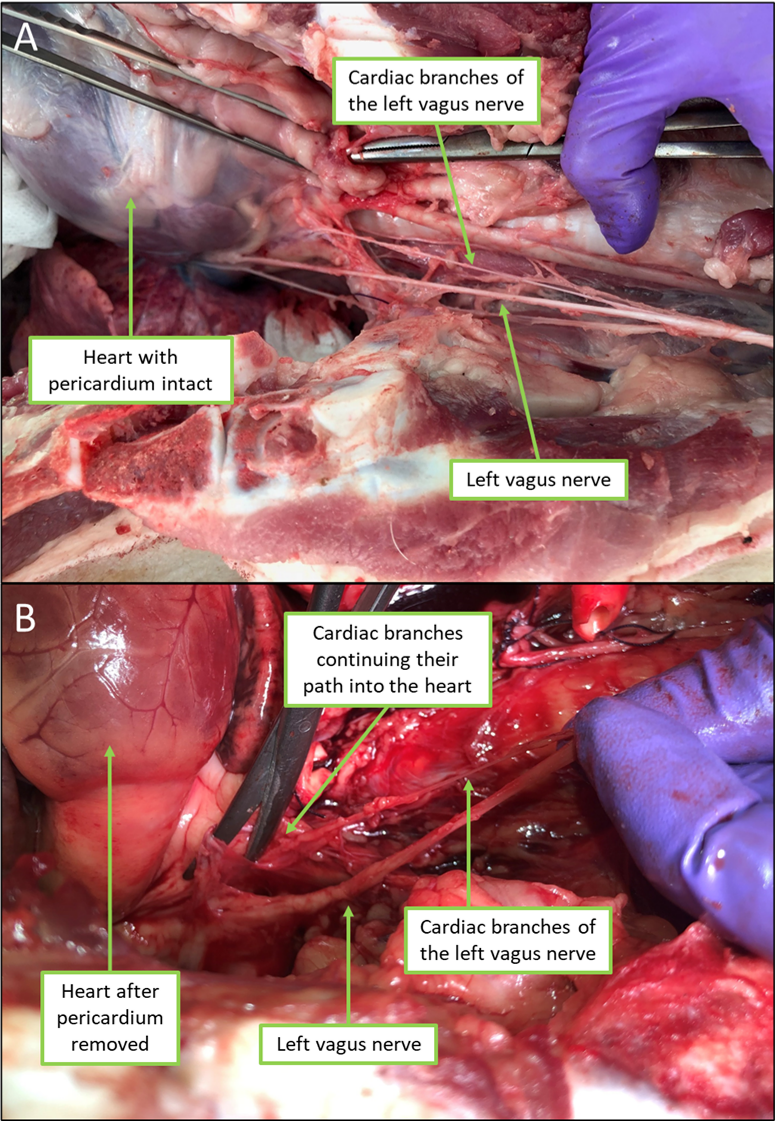

**Supplementary Figure 2. Example of vagal branch identification and dissection.** An example of dissection of cardiac branches of the left vagus nerve by tracing the branch to the organ to confirm branch type prior to labeling and cutting. The vagus nerve is initially identified and then its distal course is followed through the animal whilst dissecting any branches identified along its length. The branch is then dissected back towards its entry into the organ. Here, the branch is initially identified and the surrounding tissue dissected to visualize the path of the branch, which goes towards the heart (A). The pericardium of the heart is then removed and the branches followed further to confirm entry into the heart (B) and the branches subsequently labeled appropriately.

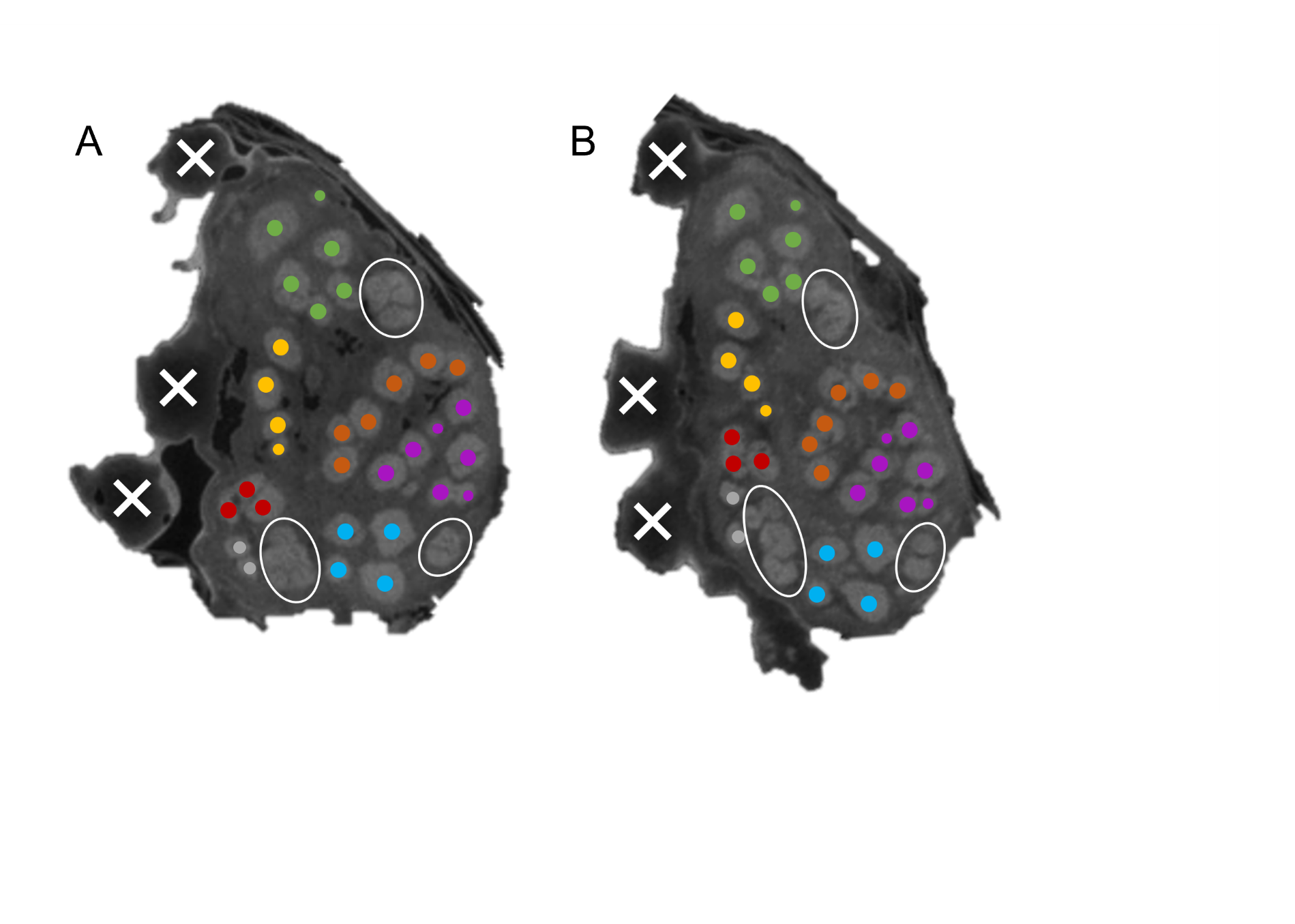

**Supplementary Figure 3. Cross sections from neighboring cut segments of nerve.** The last whole cross section visible in the last scan of a segment of nerve (**A**) is matched up to the first whole cross section visible in the first scan of the next segment of nerve (**B**). Suture landmarks were used to orient the nerve initially – shown here with white crosses. The number of suture landmarks differs between the cut regions to ensure the correct neighboring segments were matched up (an additional measure to the nerve segments being arranged in the correct order). The size, number, position and pattern of fascicles were matched up between the two cross-sections – shown here in various color groupings for ease of visualization. The three areas encircled in white here depict fascicles that were in the process of merging or splitting across this cut region; however, with process of elimination (with the other fascicles identified as matching) and viewing the scans till the respective ends, it can be deduced that these fascicles follow on from one another.

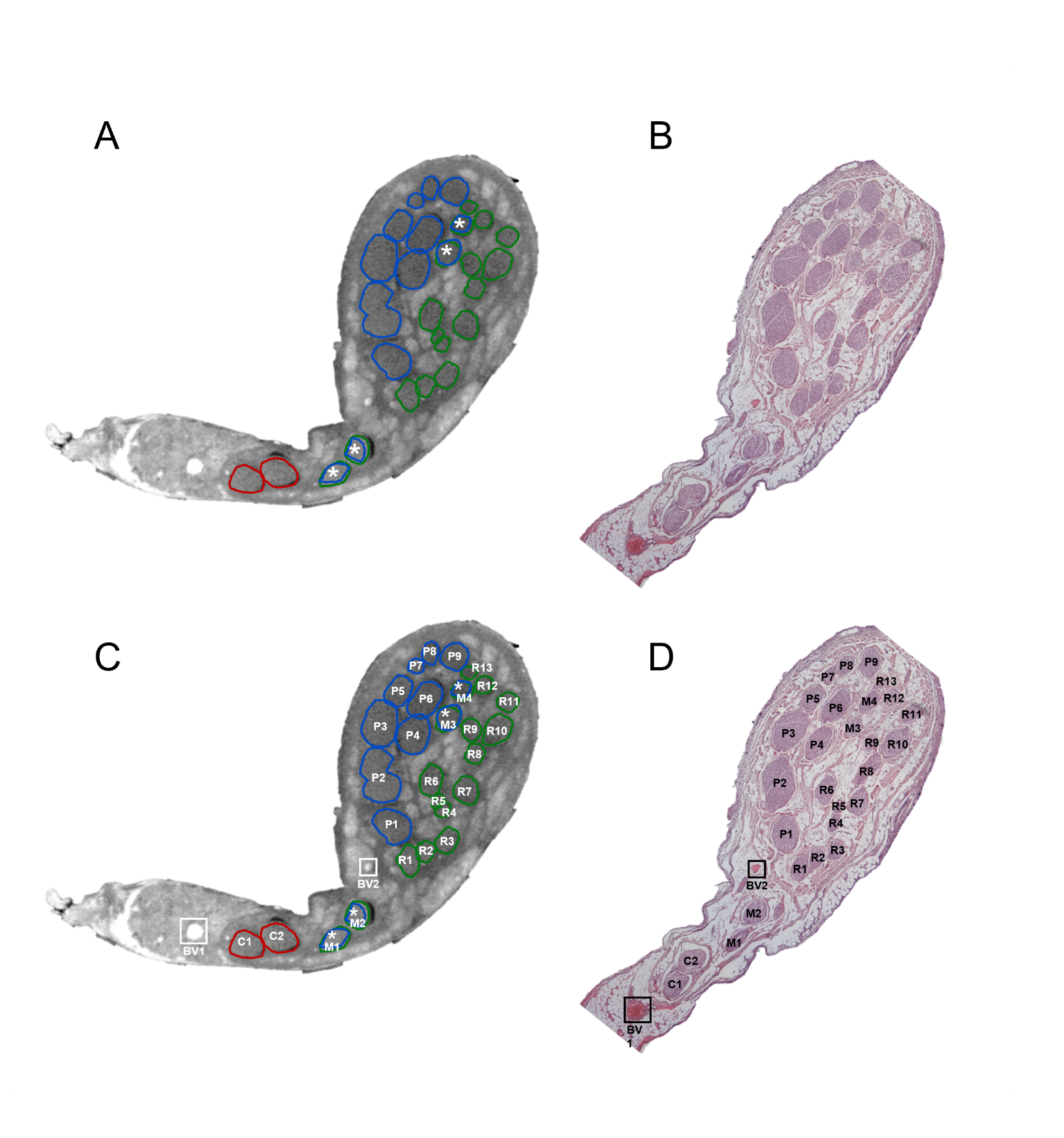

**Supplementary Figure 4. Histology validation of microCT example 1.** (A) MicroCT cross section (B) H&E histology cross section (C) MicroCT with fascicles and blood vessels labeled; (D) Histology cross section with fascicles and blood vessels labeled. Red = cardiac, blue = pulmonary, green = recurrent laryngeal, asterisk = merged fascicles between pulmonary and recurrent laryngeal, C = cardiac, P = pulmonary, R = recurrent laryngeal, M = merged fascicle, BV = blood vessel. See supplemental video 1 for visualization of the fascicles in the cross sections of the nerve.

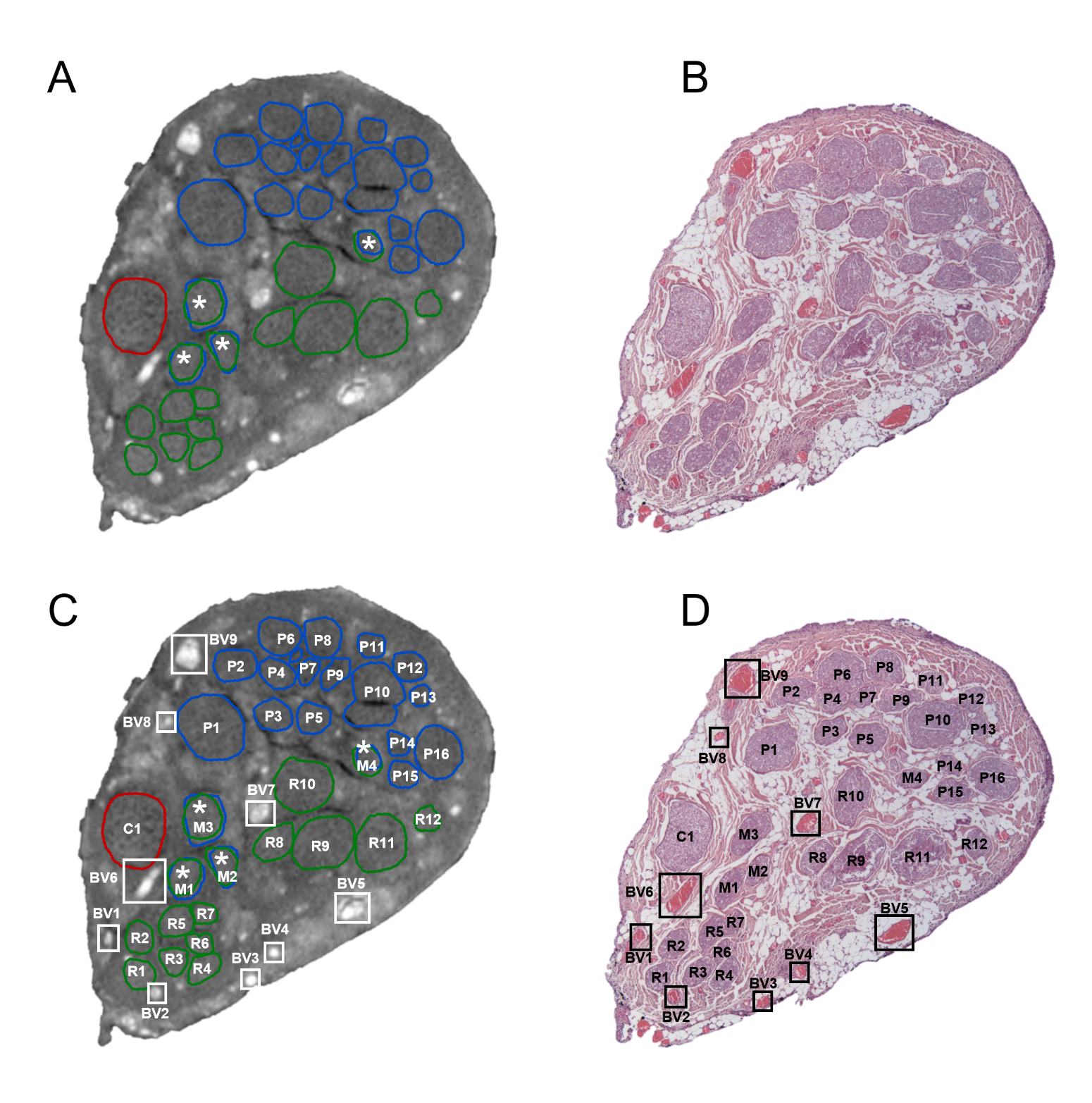

**Supplementary Figure 5. Histology validation of microCT example 2.** (A) MicroCT cross section (B) H&E histology cross section (C) MicroCT with fascicles and blood vessels labeled; (D) Histology cross section with fascicles and blood vessels labeled. Red = cardiac, blue = pulmonary, green = recurrent laryngeal, asterisk = merged fascicles between pulmonary and recurrent laryngeal, C = cardiac, P = pulmonary, R = recurrent laryngeal, M = merged fascicle, BV = blood vessel. See supplemental video 1 for visualization of the fascicles in the cross sections of the nerve.

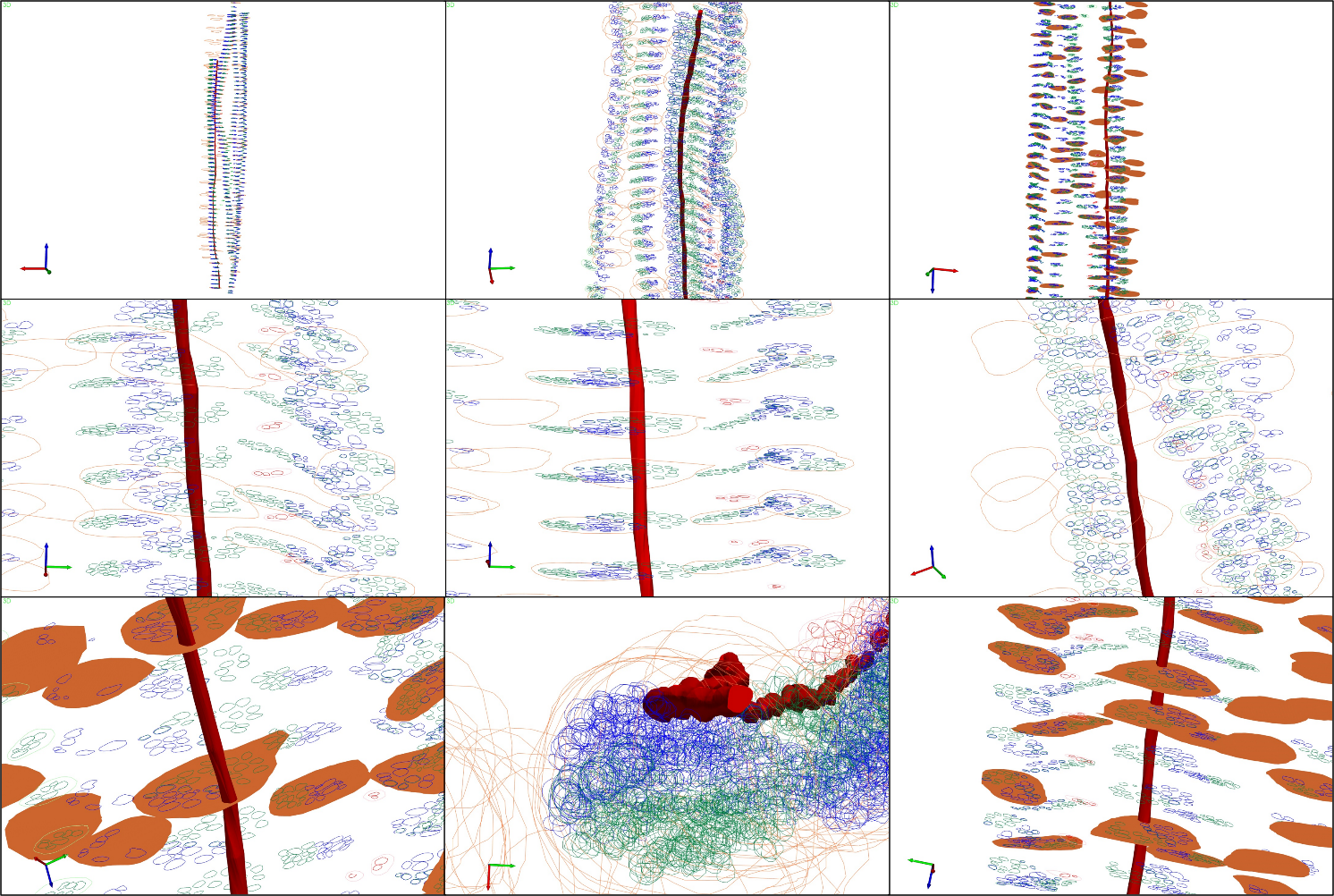

**Supplementary Figure 6. Cardiac fascicles remaining separate from the rest of the vagal fascicles.** An example of a vagus nerve with the fascicles of the cardiac, pulmonary and recurrent laryngeal branches segmented and contoured throughout the length of the nerve (the vagus nerve is in 4cm segments). The cardiac fascicles in a segment of nerve have been 3D shelled and are visualized here with the rest of the contours showing the path of the cardiac fascicle(s) through the cervical nerve without any merger with the other vagal fascicles. Red = cardiac, blue = pulmonary, green = recurrent laryngeal, orange = vagus nerve. See supplemental video 3.

#### 2.2 Supplementary Tables

Supplementary Table 1

**Selective Stimulation – Parameters and Effects**. List of neuromodulation effects achieved on the three branches of the left vagus for each animal. Effect on breathing and cardiac function is reported as beats per minute and percentage variation from baseline. Effect on larynx contraction is reported as RMS signal amplitude and relative variation from baseline. Stimulation parameters (pulse amplitude, width, repetition frequency, on/off time) are reported for the most selective iteration of stimulation on each animal and branch. Results are reported as electrode pairs with most localized effect, out of 14 radially organized pairs.

| **Recording** | | **Neuromodulation Effect** | | | **Stimulation Pulse Parameters** | | | | **Result** |
| --- | --- | --- | --- | --- | --- | --- | --- | --- | --- |
| **Pig #** | **Effect Type** | **Baseline [bpm]** | **Effect on best pair [bpm]** | **Effect [%]** | **Freq (Hz)** | **Amp (uA)** | **Width (µS)** | **On/Off time (s)** | **Active Pairs** |
| 1 | Pulmonary | 10.5 | 5.5 | -47.62 | 20 | 300 | 50 | 15 | 01 to 07 |
| 2 |  | 8.5 | 5 | -41.18 | 20 | 400 | 50 | 15 | 4, 6, 7 |
| 3 |  | 13.5 | 11 | -18.52 | 20 | 220 | 50 | 30 | 7, 8 |
| 4 |  | 8.5 | 4.5 | -47.06 | 20 | 500 | 50 | 15 | 8, 9 |
| **Pig #** | **Effect Type** | **Baseline [bpm]** | **Stim on best pair [bpm]** | **Effect [%]** | **Freq (Hz)** | **Amp (uA)** | **Width (µS)** | **On/Off time (s)** | **Active Pairs** |
| 1 | Cardiac | 87 | 82 | -5.75 | 20 | 1100 | 1000 | 30 | 1,2,3 |
| 2 |  | 65 | 55 | -15.38 | 20 | 2000 | 1000 | 15 | 1, 2, 10, 14 |
| 3 |  | 91 | 80 | -12.09 | 20 | 250 | 1000 | 30 | 4, 5, 6 |
| 4 |  | 96 | 89 | -7.29 | 20 | 800 | 1000 | 30 | 5,6,8,9 |
| **Pig #** | **Effect Type** | **Baseline [-]** | **Stim on best pair [-]** | **Effect [-]** | **Freq (Hz)** | **Amp (uA)** | **Width (µS)** | **On/Off time (s)** | **Active Pairs** |
| 1 | Laryngeal | 8.5 | 2144 | 251.24 | 20 | 150 | 50 | 5 | 8, 9 10 |
| 2 |  | 500 | 7744 | 14.49 | 20 | 250 | 50 | 5 | 9, 10, 11 |
| 3 |  | 300 | 18000 | 59.00 | 20 | 150 | 50 | 5 | 1,2,3,12,13,14 |
| 4 |  | 150 | 8180 | 53.53 | 20 | 400 | 50 | 5 | 2, 8 to 14 |

Supplementary Table 2

**EIT – Trace selection parameters, number of selected traces and resulting SNR values.** List of numerical parameters adopted for filtering δV and δV-RMS traces. Parameters were adapted to noise levels of individual recordings for laryngeal EIT. Fixed values were used for spontaneous EIT of pulmonary function. Measurement-injection pairs selected for pulmonary imaging were also selected for imaging of cardiac function.

|  |  | **Filter Thresholds** | | | | **Number of traces** | | **Peak SNR** | | |
| --- | --- | --- | --- | --- | --- | --- | --- | --- | --- | --- |
| **Pig #** | **Branch** | **Noise Std [µV]** | **Mean [µV]** | **Max [µV]** | | **Original** | **Post-filtering** | **Avg Noise [µV]** | **Mean abs (δZ) at peak [µV]** | **SNR [-]** |
| 1 | Laryngeal | 1.5 | 4.0 | 4.0 | | 196 | 92 | 0.38 | 1.25 | 3.28 |
| 2 |  | 1.0 | 4.0 | 8.0 | | 196 | 133 | 0.38 | 1.96 | 5.16 |
| 3 |  | 1.0 | 2.0 | 7.0 | | 196 | 158 | 0.47 | 2.56 | 5.39 |
| 4 |  | 4.0 | 6.0 | 7.0 | | 196 | 95 | 0.84 | 1.98 | 2.37 |
| **Pig #** | **Branch** | **Max amplitude [µV]** | **Stds from mean [µV]** | **Derivative [µV/s]** | **Stds from PCA [-]** | **Original** | **Post-filtering** | **Avg Noise [µV]** | **Mean abs (δZ-RMS) at peak [µV]** | **SNR [-]** |
| 1 | Pulmonary | 0.2 | 3 | 50.00 | 3 | 182 | 114 | 2.78E-09 | 8.00E-09 | 2.88 |
| 2 |  | 0.2 | 3 | 50.00 | 3 | 196 | 158 | 2.60E-09 | 8.31E-09 | 3.19 |
| 3 |  | 0.2 | 3 | 50.00 | 3 | 168 | 114 | 2.60E-09 | 8.31E-09 | 3.20 |
| 4 |  | 0.2 | 3 | 50.00 | 3 | 168 | 116 | 2.31E-09 | 6.86E-09 | 2.97 |
| **Pig #** | **Branch** | **Max amplitude [µV]** | **Stds from mean [µV]** | **Derivative [µV/s]** | **Stds from PCA [-]** | **Original** | **Post-filtering** | **Avg Noise [µV]** | **Mean abs (δZ-RMS) at peak [µV]** | **SNR [-]** |
| 1 | Cardiac | **Same as pulmonary** | | | | | | 3.20E-09 | 9.09E-09 | 2.84 |
| 2 |  |  |  |  |  |  |  | 2.53E-09 | 8.82E-09 | 3.48 |
| 3 |  |  |  |  |  |  |  | 3.71E-09 | 1.33E-08 | 3.60 |
| 4 |  |  |  |  |  |  |  | 4.23E-09 | 9.25E-09 | 2.19 |

#### 2.3 Supplementary Videos

Supplementary Video 1

MicroCT Nerve Scan XY Example 1 – available at <https://youtu.be/Pm6Ahd1_1wA>

Supplementary Video 2

MicroCT Nerve Scan XY Example 2 – available at <https://youtu.be/OjiMKlKVhy0>

Supplementary Video 3

Cardiac Fascicles Separated from Nerve Example – available at <https://youtu.be/kKSjirWHm9E>
